# The skull of StW 573, a 3.67 Ma *Australopithecus* skeleton from Sterkfontein Caves, South Africa

**DOI:** 10.1101/483495

**Authors:** Ronald Clarke, Kathleen Kuman

## Abstract

Here we present the first full anatomical description of the 3.67 million-year-old *Australopithecus* skull StW 573 that was recovered with its skeleton from the Sterkfontein Member 2 breccia in the Silberberg Grotto. Analysis demonstrates that it is most similar in multiple key morphological characters to a group of fossils from Sterkfontein Member 4 and Makapansgat that are here distinguished morphologically as *A. prometheus*. This taxon contrasts with another group of fossils from those sites assigned to *A. africanus*. The anatomical reasons for why these groupings should not be lumped together (as is frequently done for the South African fossils) are discussed in detail. In support of this classification, we also present for the first time a palate (StW 576 from Sterkfontein Member 4) newly reconstructed by RJC, which has a uniquely complete adult dentition of an *A. africanus.* The StW 573 skull also has certain similarities with other earlier *Australopithecus* fossils in East Africa, *A. afarensis* and *A. anamensis*, which are discussed. One of its most interesting features is a pattern of very heavy anterior dental wear unlike that found in *A. africanus* but resembling that found in *A. anamensis* at 4.17 Ma. While StW 573 is the only hominid fossil in Sterkfontein Member 2, we conclude that competitive exclusion probably accounts for the synchronous and sympatric presence of two species of *Australopithecus* in the younger deposits at Makapansgat and Sterkfontein Member 4. Because the StW 573 skull is associated with a near-complete skeleton that is also described for the first time in this special issue, we are now able to use this individual to improve our understanding of more fragmentary finds in the South African fossil record of *Australopithecus*.

## 1. Introduction

The StW 573 skull (Fig. 1) of a mature adult (informally named ‘Little Foot’ by P.V. Tobias) was discovered in 1998 during the excavation of an *Australopithecus* skeleton in the solid Member 2 breccia within the Silberberg Grotto of the Sterkfontein caves (Clarke, 1998). Here we present the first detailed anatomical description. StW 573 has been dated to ˜3.67 Ma (Granger et al., 2015), which places it as a southern African contemporary of the early representatives *of A. afarensis* from eastern Africa, i.e., the 3.6 Ma fossils from Laetoli, Tanzania (Leakey et al., 1976) and Woranso Mille, Ethiopia (Haile-Selassie and Su, 2016). Further details of the excavation, the geological context of StW 573, and debates on the dating of Member 2 can be found elsewhere in this special issue and so are not covered here (Bruxelles, submitted; Clarke, submitted).

**Fig. 1.**
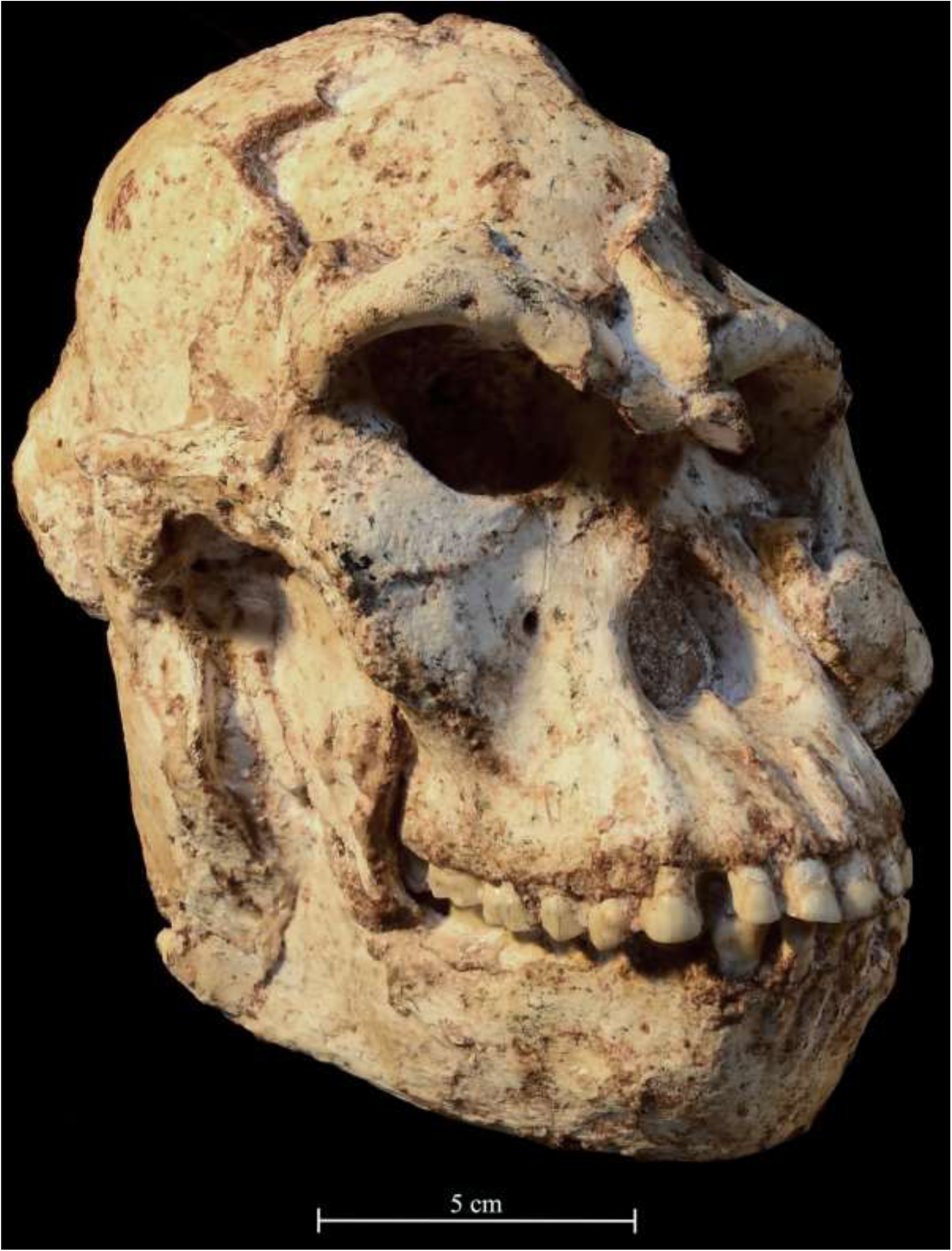
The StW 573 skull in three-quarter view. Photo by M. Lotter.

Although MicroCT scanning of the skull and other elements has been conducted since 2013 by Kris Carlson and Tea Jashashvili in the Evolutionary Studies Institute of the University of the Witwatersrand, the size of the skull, the need to remove further breccia for better results, and the lengthy segmentation process to virtually separate the bone from the very dense matrix have resulted in delays to publishing such data. As there has long been an expectation of publication, and as there is now sufficient information available on the skull to provide a meaningful description, we present what is currently discernible. Meanwhile, further study and reconstruction of the skull are ongoing and more information will be available in the near future following more physical preparation and a virtual reconstruction from MicroCT scans. The importance of this skull is: 1) its completeness, with cranium and mandible still in articulation, and the relative lack of distortion compared to the other more fragmentary crania and mandibles previously recovered from Sterkfontein; 2) the fact that it indisputably belongs with the rest of the skeleton, and 3) its great age. Its completeness means that, for the first time, it is possible to make a detailed anatomical assessment of an entire adult skull of a South African *Australopithecus*. The fact that it belongs with the skeleton also means that taxonomic placement of the skull applies equally to the postcranial bones, and thus they, in turn, provide a taxonomic reference against which other isolated postcranial remains can be compared.

## 2. Materials and Methods

The skull of StW 573 was discovered during excavation of its associated skeleton within indurated calcareous breccia in Member 2. The left side of the skull was first exposed in 1998 following removal of a 20 cm thick flowstone and then the careful working away of the breccia with an airscribe. After sustained work on excavation of the entire skeleton in situ in the cave, we were in 2010 finally able to undercut and remove the breccia block containing the skull to the Sterkfontein field laboratory. The skull was then further processed from the breccia with airscribes through October 2017, at which point it was removed to the hominid vault in the Evolutionary Studies Institute at the university, and preparation of this paper was intensified. The soft and flaky surfaces of bone were consolidated as they were exposed with Paraloid B-72 adhesive, a copolymer of ethylmethacrylate and methyl acrylate.

Original fossil cranial material used for comparisons in this study (Table 1) is curated at the Ditsong Museum in Pretoria and at the Evolutionary Studies Institute at the University of the Witwatersrand in Johannesburg. This comparative material is from the sites of Sterkfontein Caves, Swartkrans and Makapansgat. Comparisons with East African fossils were done with the assistance of casts and published data. The analytic method used in this paper focuses on traditional morphological descriptions of the specimens, with measurements provided whenever possible.

**Table 1.**
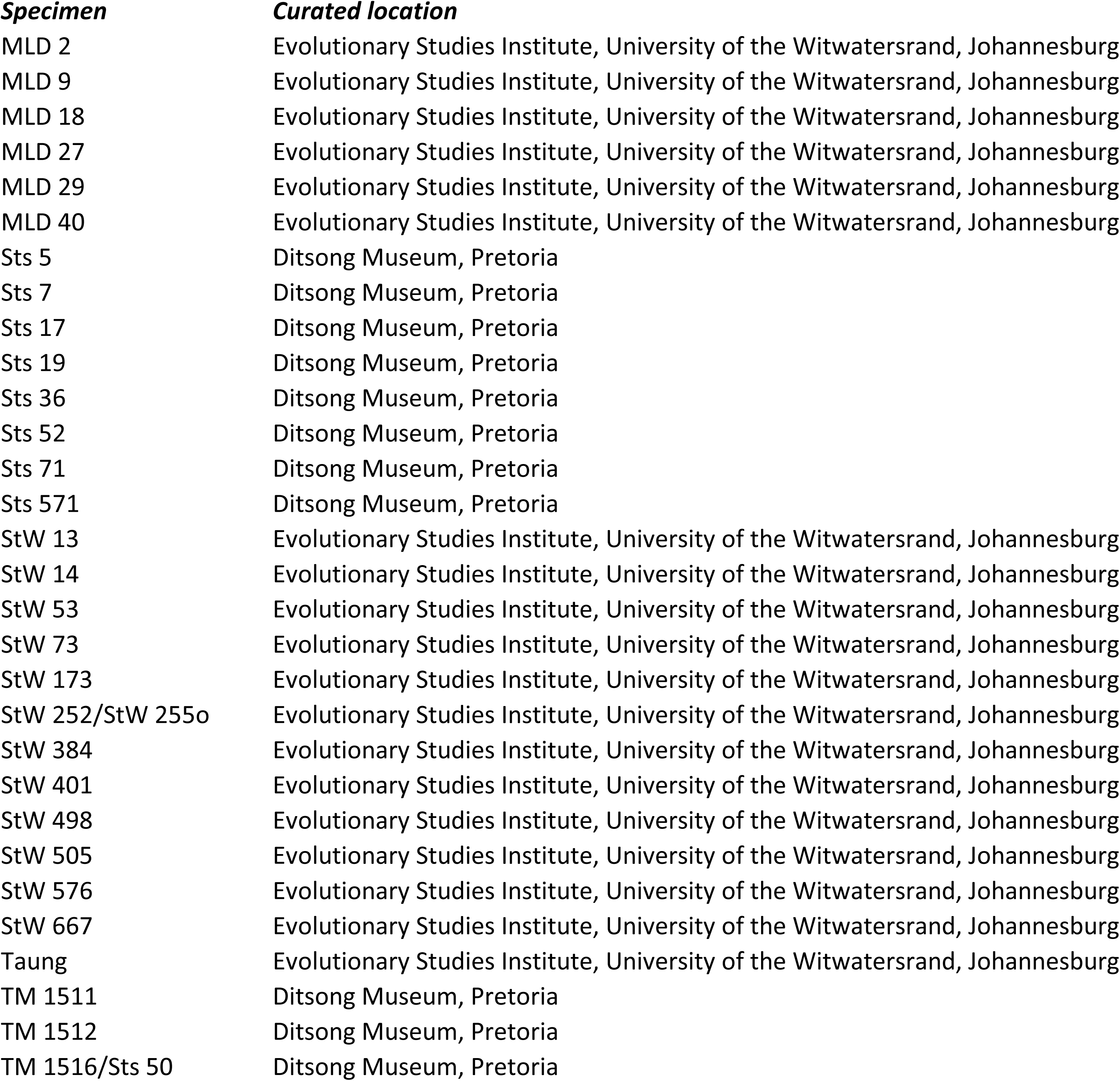
Original hominid specimens used in the comparative analysis of StW 573.

## 3. Description of StW 573

### 3.1. Condition

Although the skull (Fig. 1) has been largely cleaned of matrix, there is still some breccia filling the orbits, the nasal cavity, the auditory meati, and the left temporal fossa, as well as the right side of the endocranial cavity. The mandible is still fused by matrix in articulation and dental occlusion with the maxilla. It should be possible in the future to separate the mandible, but it will be a delicate and time-consuming operation because of the fragility of the tooth enamel which is prone to separate from the dentine, the fusing of the right condyle against the glenoid fossa, and the jamming of the right ascending ramus against the maxilla and sphenoid.

The skull of StW 573 is intact, missing only a small area of bone (20 × 22 mm) just behind the foramen magnum, and as the mandible is still in articulation, it is not yet possible to see and study the complete dental occlusal surfaces except in MicroCT scans. The bone is soft, and the surfaces tend to be flaky. Pressure within the talus slope has resulted in the displacement of the lower face upward and into the frontal and left zygomatic bones with a break through the lateral margin of the right orbit and the middle of the inferior margin of the left orbit. Upward displacement on the right is by about 16 mm and on the left by about 23 mm. The upward displacement can be seen in the midsagittal section of the skull shown in Fig. 2. The supraglabellar region is badly fragmented and buckled and glabella is projecting anterior to the upper nasal region by about 14.5 mm. The region of the left temporal line on the frontal is also buckled, and on the right side the frontal squame has been pushed downward by about 6 mm at the coronal suture. On the left side there is a break and downward displacement of the zygomatic arch just anterior to the glenoid. The anterior face of the left zygomatic bone below the orbit is cracked, with some slight displacement of part of the anterior surface. On the left side, the mandibular ascending ramus shows some cracking, with slight displacement.

**Fig. 2.**
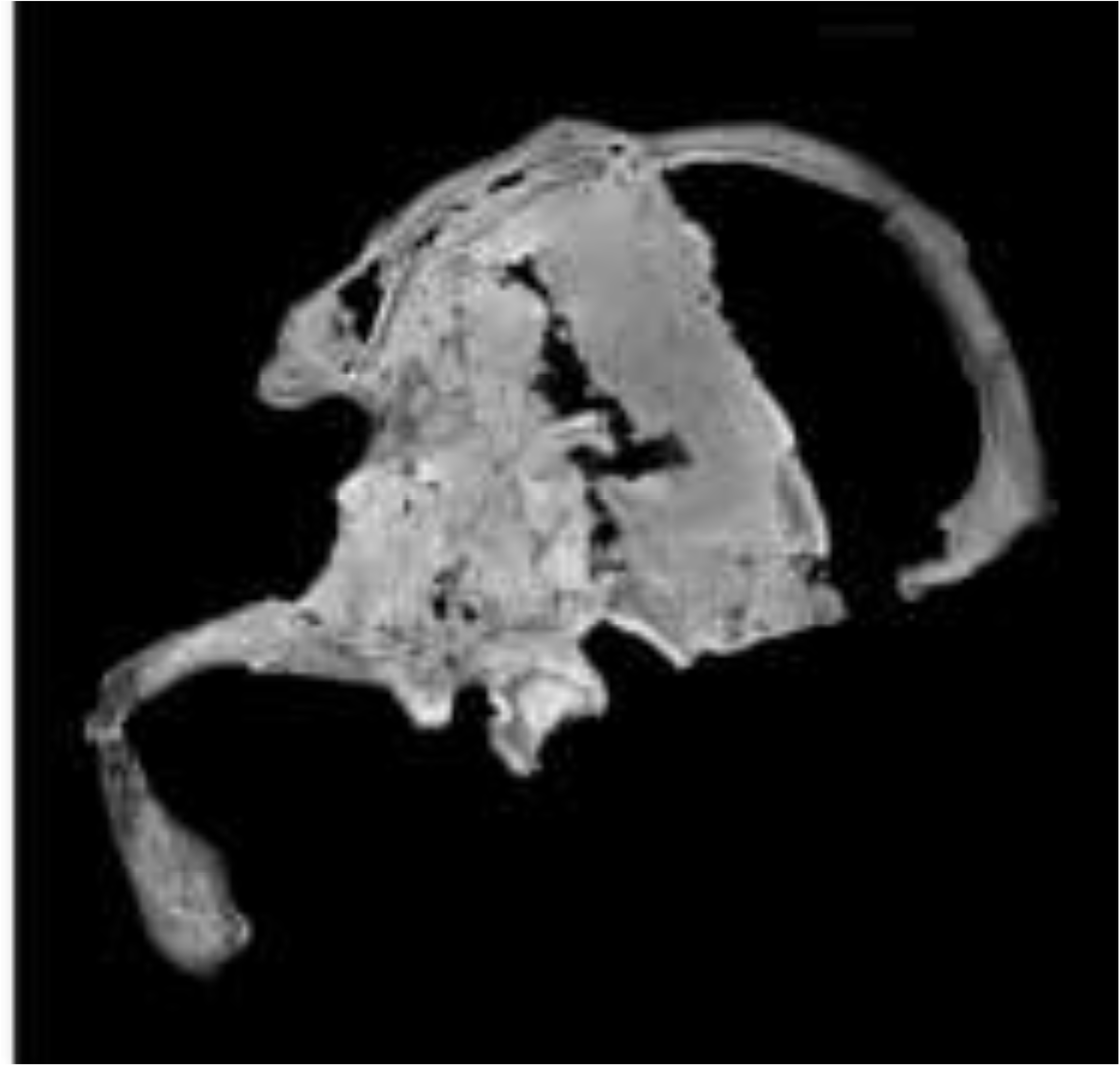
MicroCT scan of StW 573 showing the mid-sagittal section. It shows the braincase partially filled with breccia in the centre of the image. Note the calcite coatings on breccia and bone. The face has been pushed upwards and backward behind the frontal sinus. Image courtesy of Kris Carlson.

If the skull is viewed from the right lateral aspect, it is apparent that the undistorted lower and mid-face has been rotated anti-clockwise around the pivot of the temperomandibular joint, causing a fracture through the right lateral orbital margin such that the frontal process of the zygomatic passes behind the lateral end of the supraorbital margin. Here the upward displacement of the frontal process relative to the supralateral orbital margin is 18.5 mm. Centrally, the upper part of the nasal bones has been pushed behind the broken infraglabella region (Fig. 2), and on the left side there is a break running vertically from the center of the infraorbital margin to the base of the zygomatic process of the maxilla. There is a calcite-filled crack running transversely across the cranial base anterior to the foramen magnum, and there is a small area of breakage and displacement at the back of the left palate. The back of the right side of the palate is partly obscured by the atlas vertebra, which had been flipped over from its articulation with the condyles and is now closely cemented by breccia to the back of the maxilla. A small area of bone is missing from behind the posterior margin of the foramen magnum, and the adjacent surface on the left shows depressed fracturing. When the skull is viewed posteriorly, the mandible together with the back of the cranium shows an anti-clockwise skewing. There is a small depression on the left parietal caused by pressure from a dolomite block.

In the sagittal midline, the infraglabella region has been displaced posteriorly relative to the glabella by about 19.5 mm. This upward force has also broken through the right zygomatic arch anterior to the glenoid fossa, and the arch in front of that break is now situated slightly above and medial to the arch behind the break. All of this distortion has greatly reduced the size of the right temporal fossa. The mandibular ascending ramus shows vertical cracking in its central region, with some medial depression and displacement. When originally uncovered, the right clavicle was found to be jammed tightly against the length of this area (Fig. 3). It is apparent that the deformation was caused by pressure against that clavicle from above when the skull was situated with its right side downward in the talus cone.

**Fig. 3.**
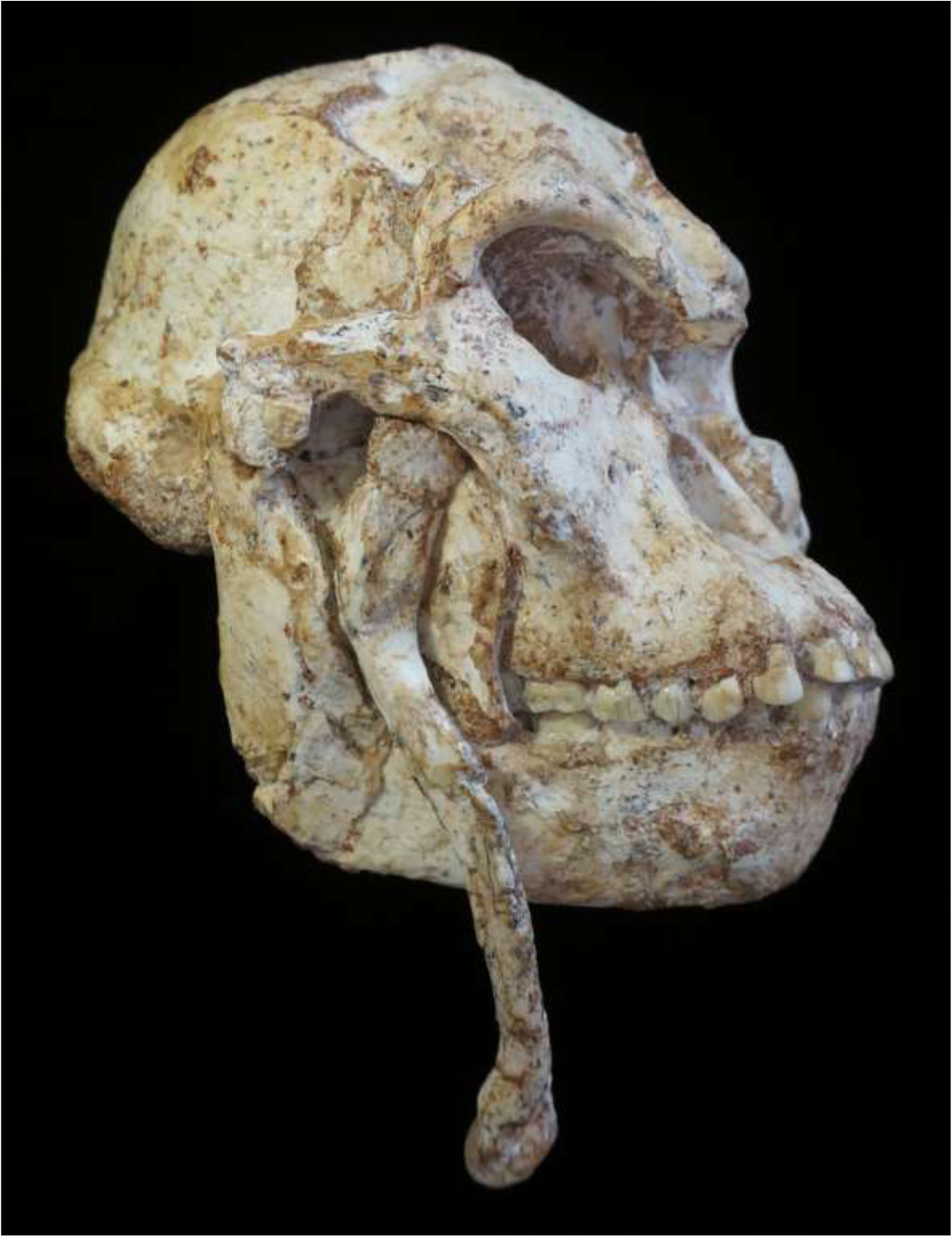
The right clavicle of StW 573 shown replaced in its original position. It was found closely cemented within a depressed area of the ascending ramus of the mandible. Photo by R.J. Clarke.

In posterior view with the base of the mandibular symphysis parallel to the nuchal plane, the right mandibular corpus and ascending ramus have a marked upward displacement relative to the left side, although there are no fractures in the corpus. The deformation was most probably associated with the upward displacement of the right and central area of the face relative to the left side. On the right, the gonial angle has suffered some fragmentation with the medial surface of the ascending ramus above it displaced somewhat laterally relative to the corpus. On the left side, the gonial angle is displaced slightly laterally and the left condyle is displaced medially relative to the corpus.

### 3.2. The cranium

#### 3.2.1 The face (Fig. 4)

The glabella is preserved at the edge of the inferiorly directed surface on the anterior end of a buckled area of the frontal squame. Posterior to glabella, this buckled squame is comprised of four fragments that, while displaced, still have good surface integrity recording a ca 15 mm broad and shallow space between the supraciliary parts of the supraorbital margin. Immediately lateral to glabella, the bone has been badly crushed upward into the frontal sinus. Superolateral to glabella, the gently concave squame gives way on each side to thickened supraorbital margins of sagittally rounded cross-section that are formed of united supraciliary and supraorbital elements. The vertical thickness of these margins is a relatively even 9 mm for most of the length, gently tapering at the lateral ends to 7.2 mm at the zygomatic suture. If allowance is made for the distortion of the squame and supraglabella region, it still seems that on both left and right sides these margins sloped gently superolaterally from the supraglabella region and then curved smoothly inferiorly at the lateral margin to meet the frontozygomatic suture. The better preserved surface of the right supraorbital margin shows clear vermiculate texture, and a trace of such texture is preserved in the centre of the left supraorbital margin. At about 22 mm lateral to glabella on each side, there is a slight supraorbital notch. That on the right is connected by a 7 mm long, faintly defined groove rising superolaterally to a small foramen on the anterior surface of the brow ridge. Posterior to the supraorbital margins and between the incurving temporal lines is a gently hollowed frontal squame that continues anteriorly to glabella and posteriorly changes to a slightly convex profile as it progresses towards bregma.

**Fig. 4.**
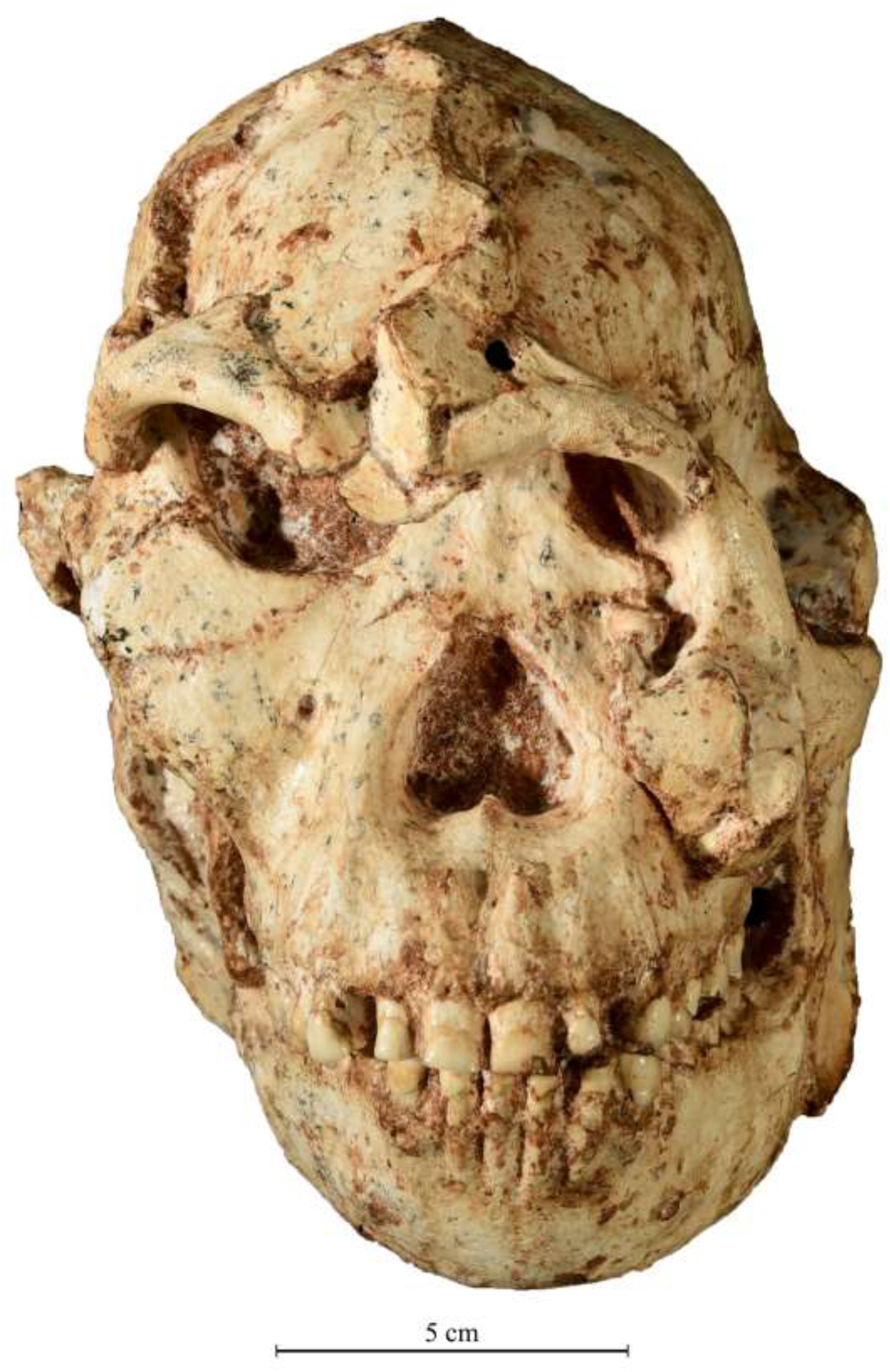
The StW 573 skull in facial view. Photo by M. Lotter.

The nasal bones are long, narrow and flat, and so well-fused to the frontal processes of the maxilla that it is not easy to determine their exact shape. They appear to be in the form of a long, narrow inverted triangle, which is flat inferiorly and forms a slight sagittal ridge superiorly. The whole upper nasal region consisting of nasal bones and adjoining frontal processes of the maxilla is flat and very broad, giving an interorbital distance (dacryon to dacryon) of 24 mm. The infraorbital region from the lateral zygomatic and extending across the lower nasal bones forms a flat surface on each side.

The complete orbital shape is not readily apparent due to the upward displacement of the lower face. The central breadth of the right orbit measures 28.5 mm, but as there is very slight lateral displacement of the lateral margin, it was possibly originally a little less. This displacement has not affected the orientation of the right frontal process of the zygomatic, which faces anteriorly. From the middle of the inferior margin of the orbit, the zygomaticomaxillary suture courses inferolaterally to the rugose masseteric insertion. This insertion is anteroposteriorly broad (12 mm at its widest) and faces laterally, tapering inferiorly to a point on the maxilla at 11 mm below the zygomaticomaxillary suture. On the left side, the well-preserved masseteric impression continues posteriorly along the inferolateral part of the zygomatic arch as a ca 11 to 10 mm broad band reaching as far as the zygomatic tubercle. The zygomatic arch is robust, being 17 mm deep anteriorly and 13 mm deep in front of the zygomatic tubercle.

From the anteroinferior tip of the right masseteric impression, the lateral margin of the zygomatic process of the maxilla descends inferomedially, presenting a gently incurved profile that terminates at the tip of the first molar roots. A small (2 mm diameter) infraorbital foramen is situated 17 mm below the inferior margin of the orbit, and a fine vascular groove ascends for 13 mm from the foramen toward the inferomedial corner of the orbit. This groove would have been for a branch of the infraorbital artery passing to the lachrymal sac. The superior margin of the foramen is 38 mm above the alveolar margin. Lateral and inferolateral to the infraorbital foramen, the cheek region is relatively flat, but inferomedially the maxillary surface curves gently anteromedially to meet the lateral margin of the nasal aperture, which presents a convex edge as it makes a tight curve into the lateral nasal wall. This provides a buttress superior to the tip of the canine root, which is situated almost at the level of the inferior margin of the nasal cavity. This root tip is 24 mm from the alveolar margin and 27 mm from the point where the enamel meets the root. The incisor roots also show very prominently and, together with the canines, present a strongly corrugated surface to the bilaterally convex nasoalveolar clivus. The tips of the large central incisor roots are 19 mm above the alveolar margin, and the tips of the lateral incisors appear to be at a similar distance, although the roots are not as prominent as are those of the canines and central incisors.

The bilaterally convex maxilloalveolar clivus with strongly corrugated surface caused by canine and incisor root eminences is a prominent feature. On the medial side of the canine juga, the concavities of the prenasal fossa are delimited anteriorly by a faint medially curving ridge emanating from the thick blunt lateral nasal margin. They are bounded posteriorly by the anterior edge formed by the nasal floor as it descends into the incisive fossa. This edge is bisected midsagittally by the inferior nasal spine that is also positioned posterior to the inferior lateral margins of the nasal cavity. This lower nasal structure has a strong similarity to that of the gorilla, from which it differs, however, by having a thicker lateral margin to the nasal cavity and the aforementioned incisor root corrugations anterior to the prenasal fossa. The frontal bone also has a general structural similarity to that of a gorilla, with the marked medially directed temporal lines delimiting an anteriorly concave frontal squame that is bounded anterolaterlly by the sagittally convex upper surface of the brow ridges.

Superior to the tips of the incisor roots, the clivus forms a slightly hollowed pre-nasal fossa which passes smoothly into the floor of the nose. A prominent inferior nasal spine is situated about 31 mm from the alveolar margin. On either side of the inferior nasal spine, the nasoalveolar clivus gives way to the depressed fossa around the incisive foramen. The nasal aperture is of an inverted heart shape measuring 24 mm in width at the level of nasospinale and 23 mm in height from nasospinale to the inferior tip of the nasal bones. Just lateral to this, the greatest height of the nasal aperture is 25 mm.

The incisors and canines are very heavily worn, but the roots and the remaining parts of the crowns show that the central incisors were relatively large with long and mesiodistally broad roots, which are 7.5 mm broad at the alveolar margin. The right central incisor crown measures 9.0 mm mesiodistally, and the left central incisor crown is also 9.0 mm. The lateral incisors are much smaller and with narrower roots. The right and left lateral incisor crowns both measure 5.8 mm mesiodistally. The right canine root mesiodistally measures 6.5 mm, and the crown is 9.5 mm. The left canine crown measures 9.8 mm mesiodistally.

The height of the clivus, i.e., the distance from nasospinale to prosthion, is 34 mm, and the greatest breadth between the outer mesial corners of the canines at the alveolar margin is 38 mm. There is a large diastema 5.6 mm wide at the alveolar margin between lateral incisor and canine on both sides.

In left lateral view (Fig. 5), the convex premaxilla with procumbent incisors is well displayed, with the 5.6 mm wide diastema between lateral incisor and canine. In right lateral view (Fig. 6), the undeformed, intact lower right half of the face is displayed from the middle of the lateral margin of the orbit and the infraglabella region downwards. From the anteroposteriorly thick (ca 15 mm) root of the zygomatic process of the maxilla, situated about 12 mm above the alveolar margin of the first and second upper molars, the anterior surface of the zygomatic process sweeps upward and only slightly backward to the orbit. Thus in this view it obscures the flat nasal bones. Anterior to the zygomatic process is a slight, shallow sulcus running inferiorly from the infraorbital foramen and forming an indistinct boundary to the low mediolateral convexity of the canine jugum. There is no marked canine fossa.

**Fig. 5.**
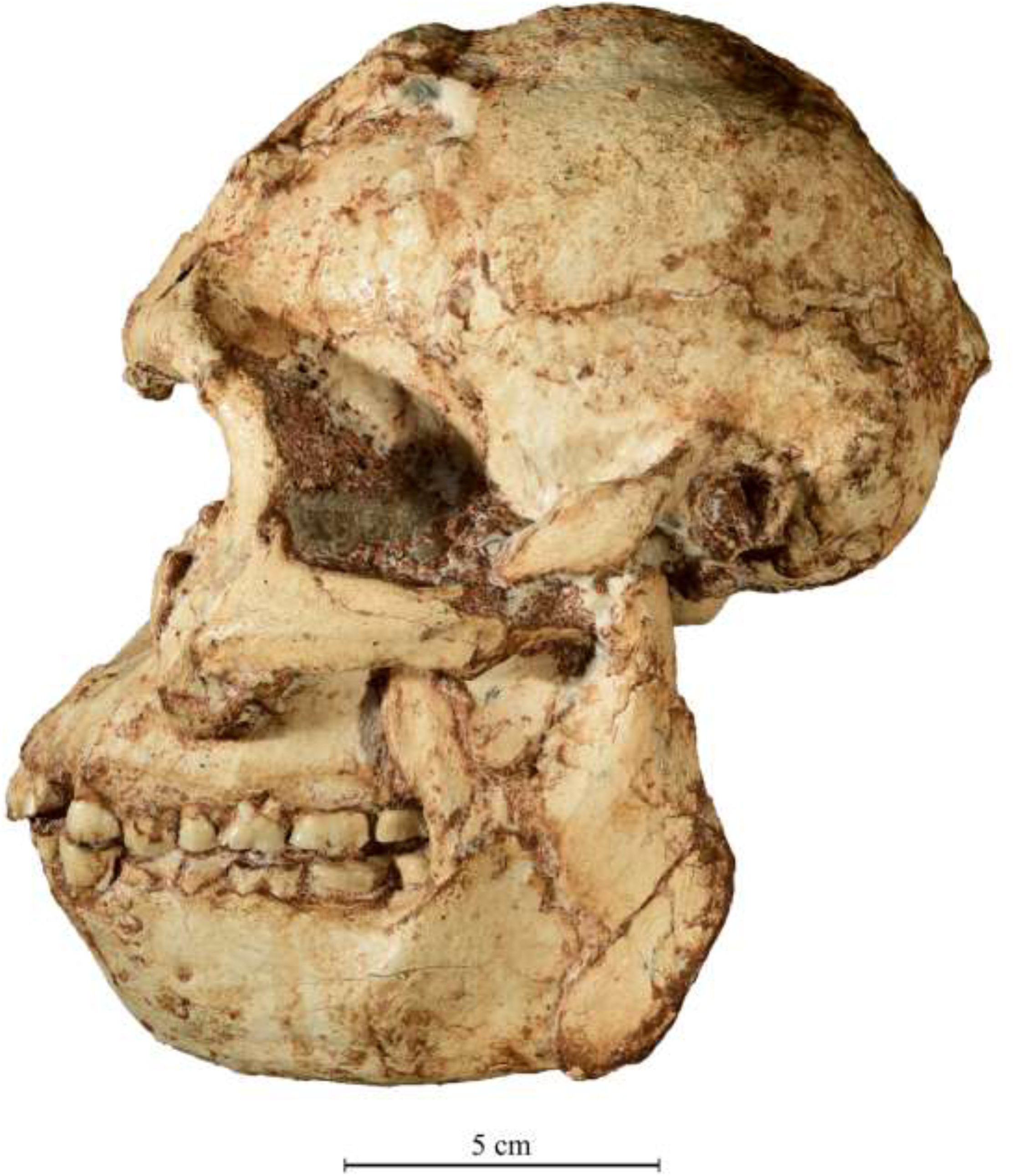
The StW 573 skull in left lateral view. Photo by M. Lotter.

**Fig 6.**
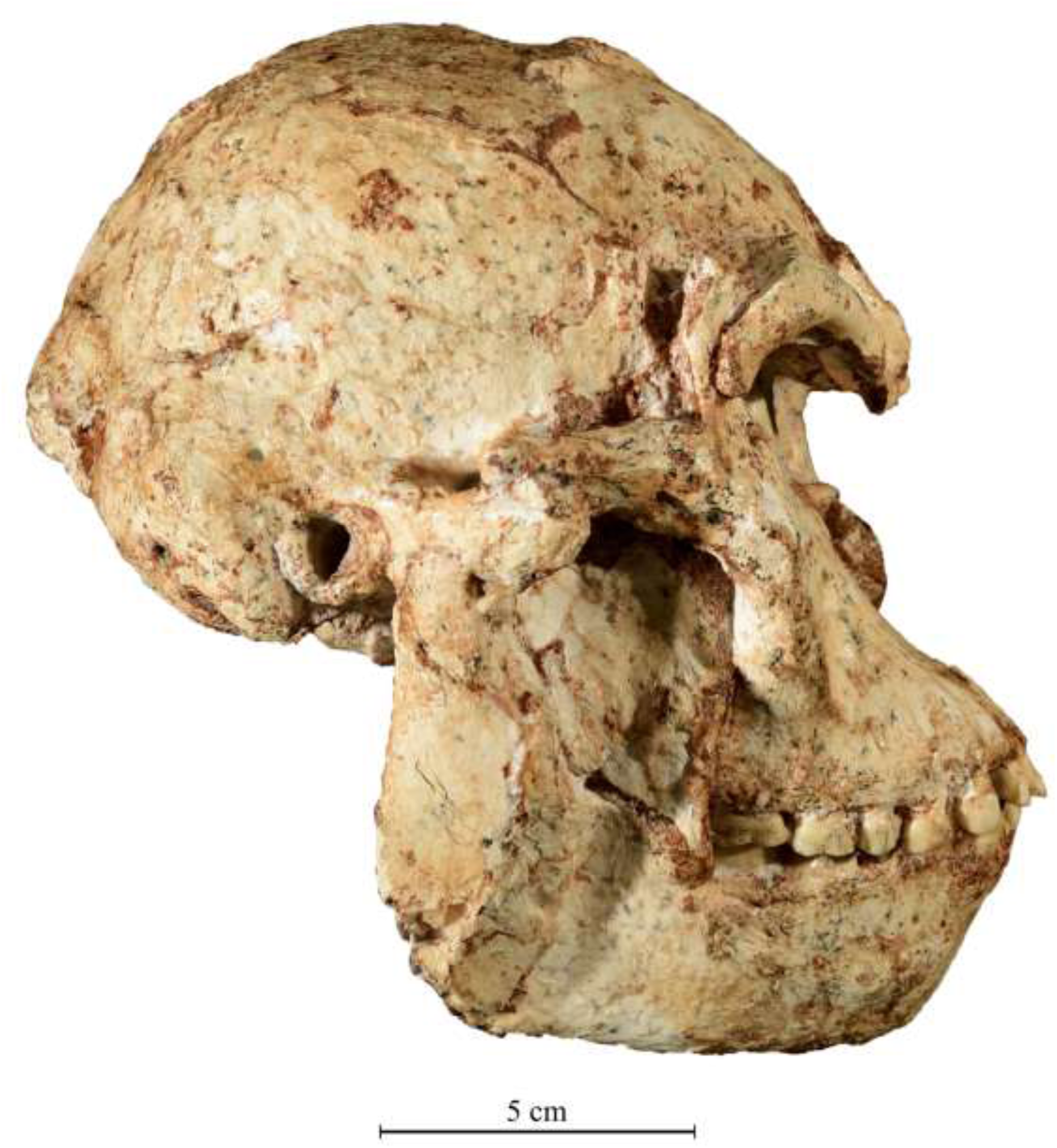
The StW 573 skull in right lateral view. Photo by M. Lotter.

From the visible infraglabella region, and with the occlusal plane horizontal, the mid-sagittal profile drops vertically to the level of the inferior orbital margin and then continues gently to an anteroinferior slope that makes an angle of ca 45°with the occlusal plane. The premaxilla is well defined, projecting anterior to the canine jugum, from which it is separated by a marked sulcus and the diastema. From a point 12 mm below the zygomaticomaxillary suture, a pronounced laterally facing masseteric impression becomes wider superiorly (11 mm), but as the zygomatic arch has been slightly eroded, the rest of the impression is not preserved.

### 3.3. The braincase (Figs. 5-8)

The frontal bone has a general structural similarity to that of a gorilla, with the marked medially directed temporal lines delimiting an anteriorly concave frontal squame that is bounded anterolaterlly by the sagittally convex upper surface of the brow ridges. The temporal lines converge at bregma to form a low sagittal crest that, midway along the sagittal midline, is formed by a bipartate crest divided by a narrow groove for 24 mm before converting to a single crest, which ends at about 11 mm anterior to lambda.

The well-preserved left temporal fossa is bounded laterally by a sagittally straight zygomatic arch that, at the lateral end of the glenoid fossa, makes an abrupt mediolateral bend to meet the temporal squame above porion. Anterior to this point is a shelf extending anteriorly for about 20 mm. The exact measurement cannot be determined as it is still obscured by matrix, but the anterior breadth is about 15 mm. There is no supraporionic crest but instead there is a relatively flat area extending posterosuperiorly from porion to the mastoid crest.

Viewed from above (Fig. 8), the braincase is ovoid in form, measuring about 151 mm from glabella to inion and 93 mm across the widest part of the parietals. It is framed, in this view, by the maxillary clivus projecting anteriorly, the zygomatic arches laterally, and the inflated mastoid crests posterolaterally. The upward displacement of the lower face has brought the face out of correct alignment with the sagittal midline of the braincase which, in this view, now aligns with the right canine jugum. Other breakage and displacements seen in this view are: 1) the broken-through right frontal process of the zygomatic pushed into the temporal fossa; 2) the broken right posterior zygomatic arch; 3) the vertical break through the left malar; 4) the break through the left zygomatic arch; 5) the sagittally buckled left frontal squame; and 6) the downward displacement of the right frontal squame relative to the parietal.

**Fig. 8.**
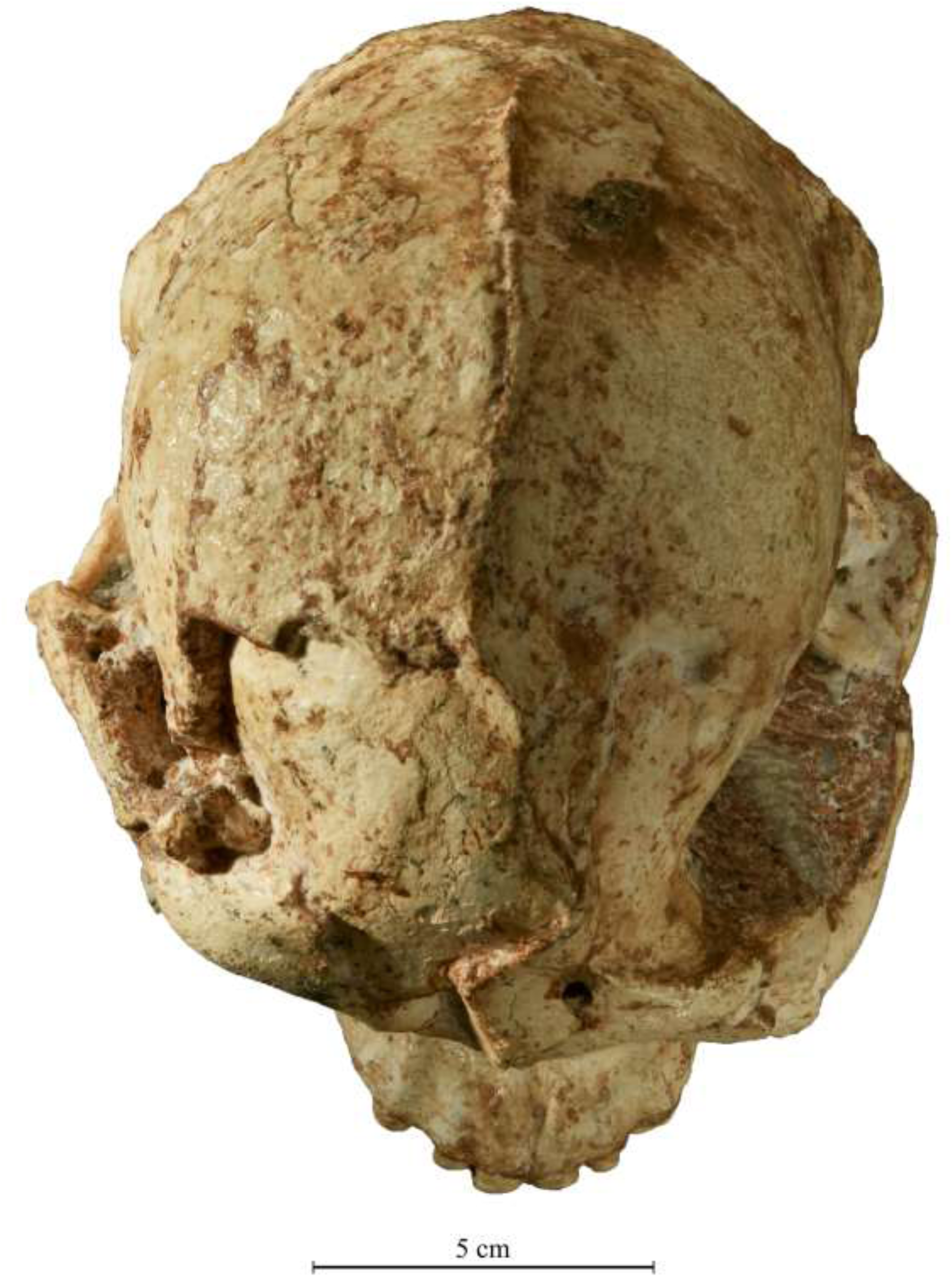
The StW 573 skull in superior view. Photo by M. Lotter.

Viewed from the left side (Fig. 5), the frontal profile behind glabella shows a slightly concave squame bounded anterolaterally by a moderately thickened (8.7 mm) supraorbital margin. That margin, of rounded anterior cross-section, is itself bounded posteriorly by the sharply medially directed temporal line that sweeps inward and upward to bregma, which is positioned on a chord 77 mm behind glabella. Posterolateral to this temporal line is a capacious temporal fossa measuring 38.5 mm at its widest, from the sphenoid to the superior margin of the zygomatic arch. The zygomatic arch is deep and robust, measuring 18.5 mm in depth from the superior junction of the arch with the frontal process of the zygomatic down to the inferior margin of the arch. At a point midway along the arch, the depth measurement is 13 mm. Inferolaterally there is a marked impression for the masseter, as described in the facial aspect. There is a ca 5 mm wide break in the arch, just in front of the supraglenoid region. The break runs inferoposteriorly, widening to about 7 mm, and was apparently caused by the wrenching and rotating upward of the lower face. The zygomatic arch anterior to the glenoid fossa measures 14 mm in depth. The length of the arch from its junction with the frontal process to the posterior root is about 63 mm when 5 mm is subtracted to account for the break and gap in the arch. The arch meets the temporal squame over the centre of the superior margin of the external auditory meatus and marks the back of a 16 mm broad supraglenoid shelf. The shelf does not continue over the ear region but instead there is a 15 mm anteroposteriorly wide, almost flat surface of the temporal squame that curves gently laterally to meet the mastoid crest.

The external auditory meatus is circular, with internal dimensions of 9.5 mm anteroposteriorly and 10.5 mm superoinferiorly. Anterior to the meatus is a very small postglenoid process. Behind the meatus is a small mastoid process that is inflated superposteriorly to form a prominent lateral swelling of the mastoid. Behind and below this swelling can be seen a broad, flat area that meets the prominent inflated occipitomastoid crest, which is visible below the level of the mastoid tip. Posterosuperiorly, the occipitomastoid crest curves strongly upward to meet and fuse with the prominent superior nuchal crest that terminates, in the midsagittal plane, in a marked and posteroinferiorly projecting process that forms inion. Between the anterolateral face of the occipitomastoid crest and the medial face of the mastoid process is an 11 mm wide well-defined digastric groove that terminates at about 18.5 mm posterior to the mastoid tip.

The nuchal plane faces posteroinferiorly. From inion, the superior profile of the braincase makes a gentle superoanterior arc to the vertex, which is situated superomedial to the glenoid fossa, and then curves on a gentle slope anteroinferiorly to glabella. A low sagittal crest occupies almost the entire length of the parietals. In the centre of the parietal bone is a slightly inflated parietal boss. The junction of the temporal squame with the parietals forms an almost horizontal line at about 26 mm above porion and 43 mm above the mastoid tip. This is because a small superior part of the temporal squame is missing. Therefore it does not match the undamaged right temporal squame which forms a straight line superoanteriorally until ca 35 mm above porion and 55 mm above the mastoid tip where it angles inferoposteriorly towards the supramastoid region.

The right lateral side (Fig. 6) shows the same morphological features as those noted for the left side, with the added advantage of an unbroken and well visible zygomatic process of the maxilla.

The posterior view (Fig. 7) shows the slight deformation that was probably caused by pressure on the left side when the skull was right side down in the talus slope. If the skull is held with the sagittal crest and inion forming a vertical line, then the right side of the braincase and the right mandibular corpus are situated higher than their counterparts on the left.

**Fig. 7.**
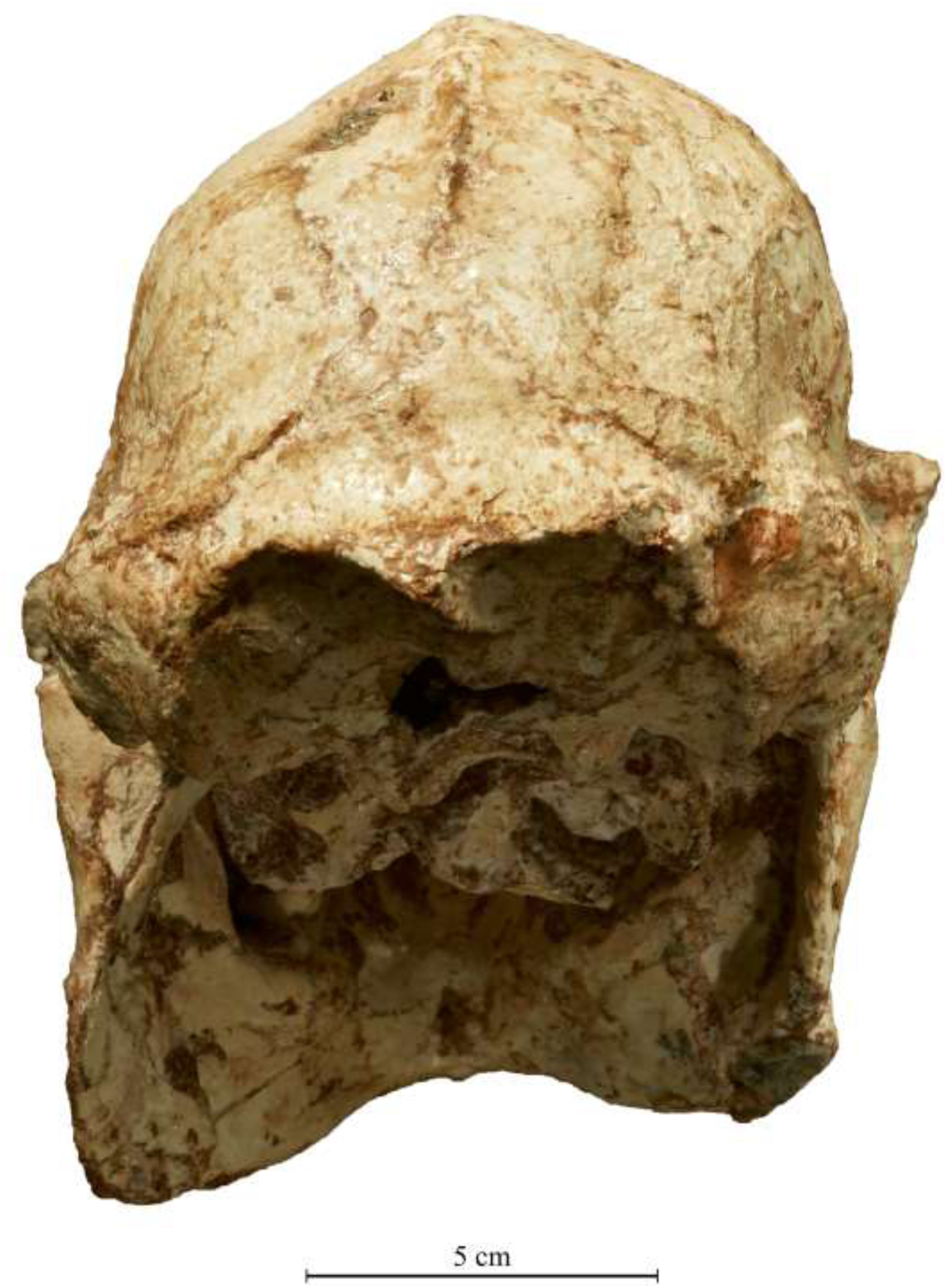
The StW 573 skull in posterior view. Photo by M. Lotter.

The braincase has the general shape of a bell with a sagittal crest at the top and almost vertical sides that flare out to the mastoid inflation at the base. At the vertex is a low sagittal crest from which the parietal surfaces slope inferolaterally, then turn downward around parietal bosses to form the almost vertical sides to the vault. The bilateral width across the bosses is 91 mm and across the temporal squame is 94 mm. If the skull is held with the nuchal plane horizontal, the parietal region of the braincase is seen to be leaning very slightly to the right. From the tops of the temporal squamae, the profile flares laterally to the mastoid crests, and then inferomedially to the mastoid tips. From the posterosuperior ends of the mastoid crests which are inflated with air cells, the temporal crests sweep superomedially to meet the sagittal crest 11 mm above lambda. From lambda, the lambdoid suture runs inferolaterally alongside but inferior to the temporal crest and meets the posterior margin of the temporal base at asterion before turning anteriorly to course along the occipitomastoid crest. The temporal and nuchal crests are not joined but are very close together (4 mm near asterion). From lambda to inion, the distance is short (36 mm), and the occipital surface between them is transversed by a marked groove running from left to right asterion. The flat and relatively smooth medioinferiorly sloping backs of the mastoids are bounded superolaterally by the convex inflated and rugose mastoid crests. The very prominent inion points inferiorly and on each side is bounded by sharply defined nuchal crests that curve superolaterally and then inferolaterally to fuse at asterion with the occipitomastoid crests. These crests project markedly inferiorly and define the lateral margins of the nuchal plane, which faces more inferiorly than posteriorly. The back of the zygomatic arch projects laterally from a point superanterior to the mastoid.

In superior view (Fig. 8) and from the frontozygomatic suture, the temporal lines sweep sharply in a posteromedial direction for about 25 mm before turning posteriorly and slightly medially to converge just behind bregma and form a low sagittal crest. Midway along its length, the crest is bipartate and divided by a narrow groove for 24 mm before converting to a single crest, which ends at about 11 mm anterior to lambda. The distance between the temporal lines at about 30 mm behind glabella is 35 mm.

The well-preserved left temporal fossa is bounded laterally by a sagittally straight zygomatic arch that at the lateral end of the glenoid fossa makes an abrupt mediolateral bend to meet the temporal squame above porion. Anterior to this point is a shelf extending anteriorly for about 20 mm. The exact measurement cannot be determined as it is still obscured by matrix, but the anterior breadth is about 15 mm.

### 3.4. The palate and basicranium (Fig. 9)

The anterior end of the palate is narrowed between the alveolar processes at the level of the incisive foramen (ca 30 mm) and relatively shallow with a long gradual slope from the incisive foramen to the incisor alveolar margin. The back of the palate is also narrow (ca 37 mm) between the alveolar processes at the alveolar margin. The roof of the palate is fragmented and displaced downward on the posterior left side. There is a relatively long maxillary tuberosity extending posteriorly for about 21 mm from the distal end of the third molar up to the join with the pterygoid plates. The medial pterygoid plate is narrow with only a narrow pterygoid sulcus separating it from the lateral plate. The lateral pterygoid plate is relatively broad ca 7.5 mm on its visible inferomedial face. There is a small triangular chip of bone missing on its distal margin. A calcite-filled crack extends across the cranial base anterior to the glenoid fossae. Posterior to this, the cranial base is well preserved. It measures 110 mm at its widest across the mastoids and 75 mm from the front of the foramen magnum to the inion. The foramen magnum is missing part of its posterior margin but can be estimated at about 25 mm in anteroposterior length. It measures 20 mm in greatest breadth at the back of the condyles. The distance from the back of the foramen magnum to the inion is notably long at 48.5 mm. Also notable is that the front and back of the foramen magnum fall almost on a straight line connecting them to the inion. In other words, the nuchal plane does not make a marked angle with the plane of the foramen magnum.

**Fig. 9.**
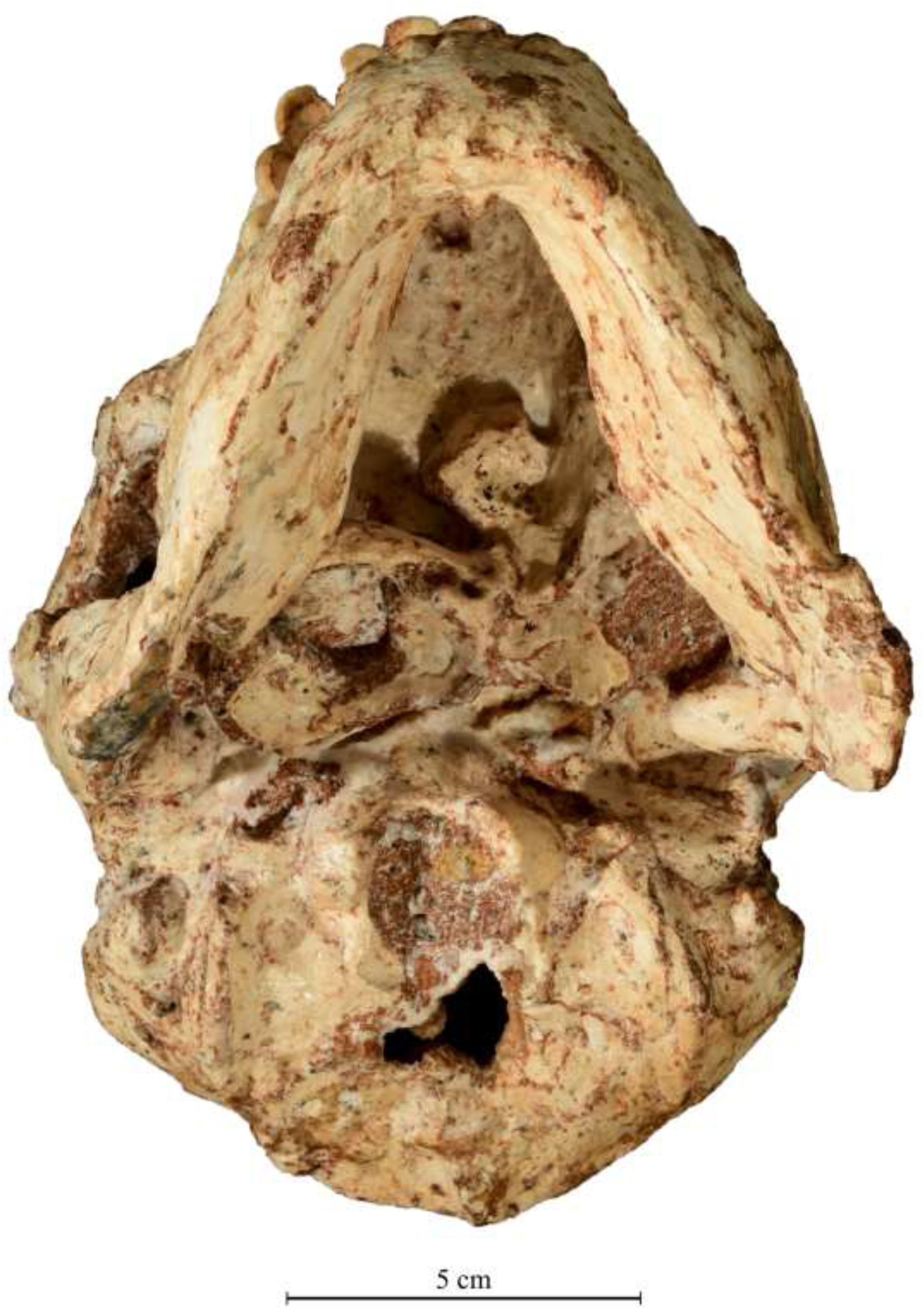
The StW 573 palate and basicranium. Photo by M. Lotter.

The better preserved left occipital condyle measures 18.5 mm in length. Lateral to the condyle, the occipital surface on each side extends horizontally for 10 mm to the occipitomastoid crest, where it then makes an abrupt upward turn for 6 mm to meet the posterior margin of the tympanic plate and the anterior end of the broad digastric fossa. On the left side, the tympanic plate is visible as an anteroposteriorly broad surface which faces anteroinferiorly in front of the digastric groove, then curves medioanteriorly and inferiorly to end as a delicate spine 12 mm anterosuperior to the lateral margin of the occipital condyle. There is no petrous crest but only a very faint flattened ridge that indicates a change in contour from the anteroinferior to the inferior surface of the plate. There is no vaginal process. The centre of the glenoid fossa is situated about 15 mm higher than the inferior margin of the tympanic plate. Posterolateral to the condyle is a marked elongate condylar fossa, separated by an anterolaterally oriented ridge from a small round fossa for insertion of the rectus capitis lateralis muscle. This fossa abuts the medial face of the prominent occipitomastoid crest. On either side, these crests dominate the topography of the nuchal plane. The bilateral distance between the summit of these crests is 61 mm anteriorly, 65 mm at the level of the mastoid tips, and 72 mm posteriorly before they curve medially towards inion. The crests are mediolaterally thick (ca 10 mm) and project well below the tips of the mastoid processes. The crest on the left is marked along its length by a 2 mm wide groove. The lateral face of the anterior half of each crest is impressed by the digastric fossa that separates the crest from the relatively small mastoid process. The tip of the process is about 5 mm above the crest on the left and slightly less on the right. From the tips of the mastoids, the mastoid crests sweep laterally and then medially to fuse with the posterior expression of the temporal crests. Inferior to these temporal crests, the occipitomastoid crests have merged posteriorly with the strongly defined inferior nuchal crests, which meet at inion. From that point, a prominent external occipital crest courses anteriorly with a slight deviation and inclination to the right side.

In this inferior view, the nuchal region appears in the form of a sharply defined circle formed by the condyles and the occipitomastoid and nuchal crests and bounded on the left and right by the flattened expanse of the posterior face of the mastoids.

### 3.5. The mandible (Table 4)

Although the mandible is still locked in articulation with the cranium with the upper and lower teeth occluded, there is a slight gap separating the occlusal planes of the upper and lower tooth rows on the left side (Fig. 5). The mandible has been slightly displaced to the left such that the midsagittal line of the cranium between the upper incisors is situated above the lateral margin of the right lower central incisor. This shift, combined with the maxillary dentition being wider than the mandibular, and the aforementioned gap on the left, means that much of the lateral occlusal surface of the upper teeth is visible. They display extremely heavy lingually sloping wear on the anterior dentition, with less wear on the molars. This wear is also well seen in the mandibular teeth in the MicroCT scans (Fig. 10; and see section on dentition below). In facial view, the whole mandible is seen to be large and deep (42 mm at the symphysis), with high ascending rami.

**Table 4.** Comparative mandible measurements for StW 573, in mm.

|  | **StW 573** | **Sts 36** | **Sts 52** |
| --- | --- | --- | --- |
| Mandibular condyle breadth | 30.0 (left) | 21.5 (left; eroded) | 21.0 (right) |
| Greatest mandibular corpus thickness between M2 and M 3 | 24.2 (left) | 24.0 (right) | 27.3 (right) |
| Greatest mandibular height gnathion to infradentale | 42.6 | not measurable | not measurable |
| Mandibular height at left M1 | 34.6 | 35.5 | 28.4 |
| Left ascending ramus breadth | 45.0 (estimate due to cracking) | 52.0 | not present |
| Right ascending ramus breadth | 53.0 (estimate due to crushing) | ca 49.0 | 45.4 |
| Maximum ramus height: condylion superior to gonion | ca 78.0 (estimate due to cracking) | 72.5 | not measurable |
| Mandibular length: horizontal component of pogonian to gonion | 107.0 | 114.5 | ca 90.0 |
| Maximum ramus breadth across condyles | 121.5 | warped | not measurable |

**Fig. 10.**
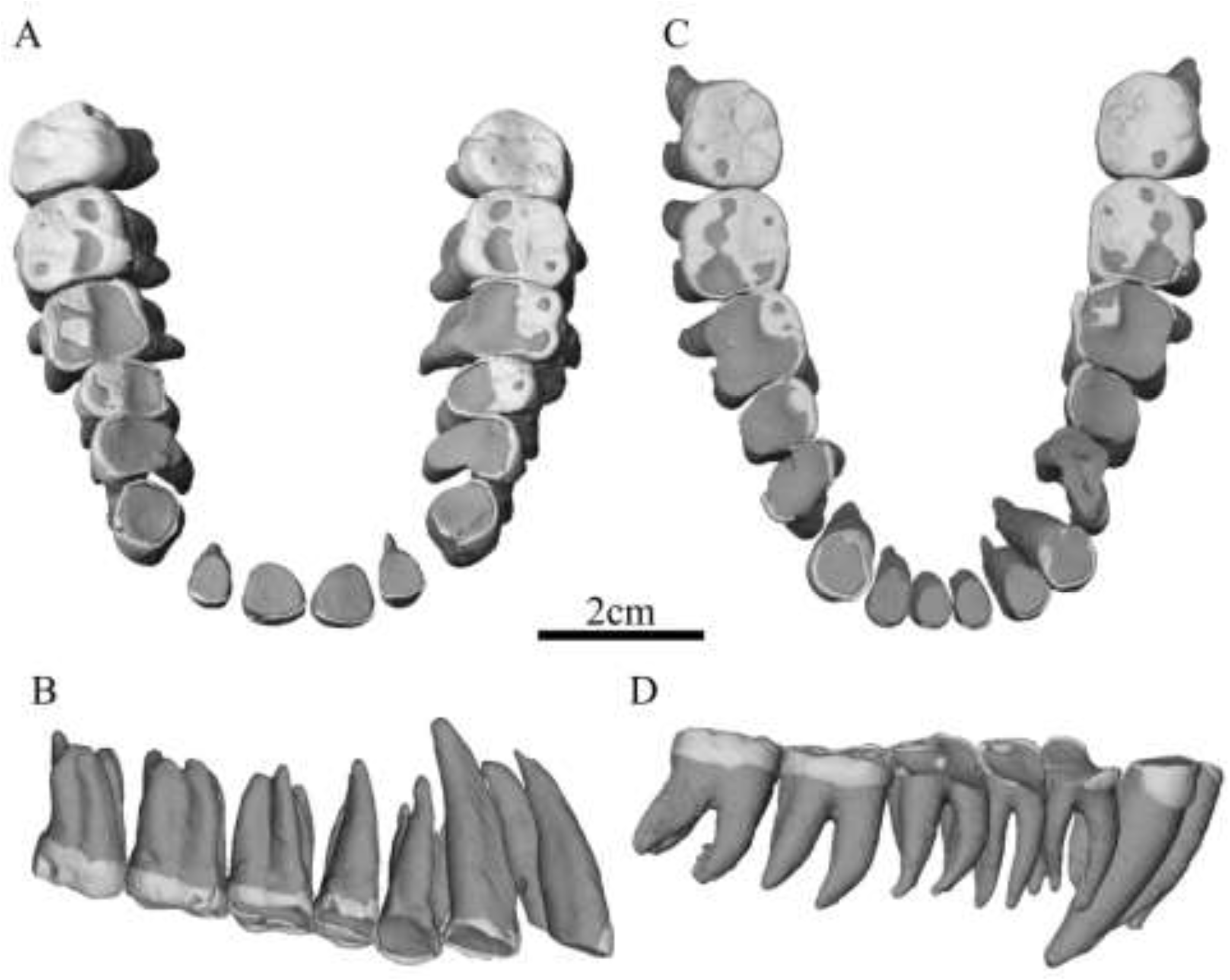
MicroCT scans of upper and lower dentition of StW 573. A) occlusal view of maxillary dentition. B) lingual view of left half of maxilla. C) occlusal view of mandibular dentition. D) buccal view of right half of mandible. Image courtesy of Tea Jashashvili.

The distance between the buccal surfaces of the upper canines is 48.5 mm, whilst that between the lower canines is 36.2 mm, showing that, in this one individual, the mandibular anterior dentition is much narrower than the upper dentition. This greater breadth of the maxilla is associated with the presence of the diastema. The roots of the lower canines form slight swellings in the mandibular surface, but the incisor roots form prominent ridges, thus giving a corrugated effect like that of the upper incisor roots.

The symphysis has a straight posteroinferiorly sloping profile, which measures 42.6 mm in depth. The depth of the corpus at M1 is 34.6 mm. The inferior margin of the corpus forms an almost straight profile but with a gentle inferior convexity that converts to a very slight concave profile at the base of the ascending ramus. The left lateral face of the mandibular body shows a slightly concave surface inferior to the P3 to a vertical depth of 18 mm, where it meets a modestly defined horizontal ridge that runs posteriorly and then superiorly to meet the oblique line in a swollen area at the junction with the anterior edge of the ascending ramus. The ridge is bounded inferiorly by an anteroposteriorly elongate shallow concavity. There are three small mental foramina. One is situated at 13 mm below the alveolar margin at the P3-P4 junction. The second is 4 mm below and 1 mm anterior to the first. It is situated at the front of the aforementioned ridge. The third foramen is also on the ridge 7.5 mm posterior to the second. The ascending ramus has several cracks accompanied by very slight lateral displacement of affected parts but it does not appear to have significantly affected the breadth or height of the ramus. The breadth of this left ascending ramus, estimated due to the cracking, is 45 mm. The maximum height of the ramus from gonion to condylion is estimated at 78 mm.

The anterior margin of the ramus presents an elongated S-shaped profile curving anteriorly at the top and distally concave in the lower part. The distal border of the ramus is almost straight from the back of the condyle to the level of the occlusal plane, where it then angles anteroinferiorly to meet the basal margin. On the lower medial surface, there are prominent pterygoid tuberosities.

The inferior view (Fig. 9) shows the great size of the mandible in conjunction with a relatively small and narrow cranial base. The base of the mandibular symphysis is 18 mm in anteroposterior thickness, just lateral to the prominent mental spine. On each side of the spine are impressions of the digastric fossae. This basal view of the mandible shows it as an equilateral triangle with the mandibular corpora forming two sides, each measuring 112 mm, and the third side being a 110 mm chord between the widely flared gonial angles at the margins of the ascending rami.

The mandibular symphysis presents a rounded bilateral profile stretching between the eminences around the mental foraminae, the distance between which is 45 mm. The corpora are, for the size of the mandible, relatively slender, with the greatest thickness half way up the corpus being 22 mm at M1 on the right and 21 mm at M1 on the left. The thickness is 24.5 mm at M2. The corpus then thins inferiorly to about 12 mm along the anterior half of the base. At a point directly inferior to the M3, the base then tapers to about 6 mm in breadth up to the gonial angle. At the anterior end of the mandibular base, the digastric fossae consist of elongate, lightly impressed marks on either side of a small anterior spine at the base of the symphysis. This spine is 5.5 mm anterior to the prominent mental spine that is situated just above the distal margin of the symphysial base. Each digastric fossa is about 7 mm broad anteriorly and tapers to a point about 24 mm distal to the symphysis.

Between the thicker part of the corpus and the base, a slightly concave area representing the subalveolar fossa can be seen anteriorly on the medial face. The left mandibular condyle is slightly twisted out of articulation with the glenoid fossa. It measures 29.3 mm in breadth. The right condyle is still in articulation but shows some damage.

The inner view of the mandible shows, from the sagittal midline anteriorly, a relatively long post-incisive plane sloping inferoposteriorly to a superior transverse torus. The torus is undercut by a well-defined genioglossal fossa. Inferior to the fossa is a pronounced mental spine, measuring about 7 mm superoinferiorly and 3.5 mm in thickness bilaterally. This spine covers the whole midsagittal depth of an inferior transverse torus, the inferior margin of which is about 4.5 mm above the lowest point of the symphysis.

The lateral halves of the mandibular condyles are visible in posterior view, as they project laterally beyond the lateral limit of the mastoids. The maximum breadth across the condyles is 123 mm.

### 3.6. Dentition (Fig. 10; Table 5)

As the jaws are still clamped together, it is not possible to see the complete occlusal surfaces, except in MicroCT scans, which have been provided by Tea Jashashvili. Her analysis from these scans will be published separately. For this paper, she has provided the occlusal and lateral views (Fig. 10). It is readily apparent that the anterior part of the dentition as far back as the first molars is very heavily worn, such that in the lower dentition the first molars retain only very small patches of occlusal enamel, whilst the premolars, canines and incisors have the occlusal enamel completely worn away. In the upper jaw, the first molars and fourth premolars have retained patches of enamel on their buccal halves, whilst the third premolars, canines and incisors have had all their occlusal enamel worn away. By contrast, in both upper and lower jaws, the second molars have retained enamel over much of their surfaces. The upper third molars are completely covered in enamel, and the lower third molars are covered in enamel except for a very small dentine exposure mesiobuccally.

**Table 5.** Grouping of morphological features for Sterkfontein and Makapansgat hominids using the best examples.

|  |  |  |  |  |
| --- | --- | --- | --- | --- |
|  | GROUP A | EXAMPLES | GROUP B | EXAMPLES |
| 1) | large face | StW 505 | smaller face | StW 53 |
| 2) | large braincase | StW 505, MLD 1 | smaller braincase | StW 53 |
| 3) | laterally expanded parietals | StW 252 | sides of braincase slope | StW 53 |
| 4) | low concave frontal squame | StW 252, StW 505 | high convex frontal squame | StW 53 |
| 5) | no metopic ridge | StW 252, StW 505 | metropic ridge | StW 53 |
| 6) | temporal lines converge medially behind brow ridge | StW 252, StW 505, Sts 71 | less convergence of temporal lines | StW 53, Taung |
| 7) | small sagittal crest on adult parietals | StW 505 | no sagittal crest on adult parietals | StW 53 |
| 8) | broad interorbital distance | StW 252, StW 505 | narrow interorbital distance | StW 53, Taung |
| 9) | flat midface | StW 252, StW 505, Sts 71 | convex midface | StW 53, Taung |
| 10) | deep malar | StW 505, Sts 71 | shallow malar | StW 53, Taung |
| 11) | robust zygomatic arch | StW 505 | more gracile zygomatic arch | StW 53, Taung |
| 12) | large masseteric attachment | StW 505, Sts 71 | reduced masseteric attachment | StW 53 |
| 13) | no supraporionic shelf | StW 505, StW 648, Sts 71 | supraporionic shelf | StW 53, Sts 5, Sts 19 |
| 14) | inferior nasal margin smoothly concave | MLD 9, StW 183, StW 498 | inferior nasal margin with transverse ridge | StW 53, StW 576, Sts 5, Sts 52 |
| 15) | procumbent upper incisors and roots | Sts 252, StW 498 | vertically oriented incisors and roots | Sts 52, StW 576, TM 1512 |
| 16) | clivus corrugated by incisor roots | StW 505 | relatively uncorrugated clivus | StW 576, StW 53 |
| 17) | inferior margin of zygomatic process runs superolaterally | StW 505 | inferior margin of zygomatic process runs more laterally | StW 53, StW 576, Sts 5 |
| 18) | large canines | TM 1516, StW 252, StW 183, StW 667 | smaller canines | StW 576 |
| 19) | large flattened upper canine roots | StW 252, StW 667 | smaller round-sectioned upper canine roots | StW 576 |
| 20) | shallow anterior palate | MLD 9, StW 252, StW 183 | deeper anterior palate | StW 53, Sts 5, MLD 6, Sts 52 |
| 21) | long rectangular palate | StW 252 | short square palate | StW 53, Sts 5 |
| 22) | upper incisor-canine diastema | StW 252, StW 183 | no upper incisor-canine diastema | TM 1512, StW 576, Sts 52 |
| 23) | large cheek teeth | StW 252, StW 183, MLD 2 | smaller cheek teeth | Sts 52, StW 53, StW 576 |
| 24) | sides of cheek teeth slope towards occlusal surface | MLD 2, StW 252, StW 183 | vertical sides to cheek teeth | Sts 52, StW 53, StW 576 |
| 25) | low bulbous cheek teeth cusps | MLD 2, StW 252, StW 183 | higher pointed cheek teeth cusps | Sts 52 |
| 26) | excessive anterior tooth wear | MLD 9, Sts 36 | more generalised tooth wear | StW 576 |
| 27) | large mandible with big teeth but relatively slender corpus | Sts 7, Sts 36, StW 384, StW 498 | small mandible with smaller teeth but thicker corpus | MLD 40, MLD 18 |
| 28) | prominent occipitomastoid crest | StW 648 | low occipitomastoid crests | StW 53, Sts 5 |

The upper dental arcade shows a very large diastema between canines and procumbent incisors, whilst the lower arcade shows a diastema between the canines and the third premolars. In the upper jaw, the central incisors are much larger than the laterals, whilst in the lower jaw the lateral incisors are much larger than the centrals. The scan of the buccal side of the lower teeth reveals the extent of sloping buccal wear on the premolars and molars, the robusticity of all the roots, and the length and great size of the canine root. The scan of the lingual side of the upper dentition shows excessive lingually sloping wear on the anterior dentition, with little wear on the third molar. The robusticity of all the roots and the great length of the incisor and canine roots are well displayed. Tooth measurements are provided in Table 2.

**Table 2.** Cranial measurements (in mm) of StW 573 and some comparative crania. Values in brackets are estimates. Abbreviations: Sts = Sterkfontein Type Site); SK = Swartkrans; OH = Olduvai Hominid); KNM-ER = Kenya National Museum, East Rudolf. Comparative measurements are from Tobias (1967) and Wood (1991).

| MEASUREMENT | StW 573 | SPECIMENS |  |  |  |  |  |  |
| --- | --- | --- | --- | --- | --- | --- | --- | --- |
|  |  | Sts |  | SK | OH | KNM-ER |  |  |
|  |  | 5 | 71 | 48 | 5 | 406 | 1470 | 1813 |
| Maximum cranial length: glabella to opisthocranium | **[151]** | 147 | 129 | 170 | 173 | 163 | 166 | 149 |
| Maximum cranial breadth: euryon to euryon | **110** | 111 | 110 |  | 142 | 144 |  | 113 |
| Bizygomatic breadth: zygion to zygion | **130** | 126 | 126 | 145 | 168 | 179 |  | 117 |
| Maximum cranial height: basion to bregma | **[104]** | 101 |  |  | 99 | 103 |  | 90 |
| Upper facial height: prosthion to nasion | **[87]** | 77 | 71 | 80 | 112 | 88 | 90 | 66 |
| Upper facial breadth: frontomale temporale to frontomale temporale | **[90]** | 94 | 92 | 107 | 115 | 115 | 114 | 100 |
| Orbital breadth: dacryon to ectoconchion | **[30]** | 34 | 34 | 33 | 40 | 39 | 41 | 33 |
| Interorbital breadth: dacryon to dacryon | **24** | 16 | 20 | 24 | 23 | 27 | 23 | 20 |
| Nasal aperture breadth: alare to alare | **27** | 26 | 24 | 29 | 32 | 28 | 27 | 23 |
| Mid-nasal aperture height: nasospinale to rhinion | **24** | 24 | 28 | 23 | 34 | 36 | 39 | 28 |

The upper left tooth row from front of canine to back of third molar measures 60 mm.

## 4. Comparative observations

### 4.1. Condition

Considering that the skull was situated beneath and among many large and small dolomite and chert blocks within the talus slope and that the braincase is only partially filled with breccia, it is fortunate indeed that the damage and deformation were not more extensive and that the dentition has been preserved intact, despite the fact that the enamel is not well-bonded to the dentine. Not only is this the most complete skull of any *Australopithecus* yet discovered, but it has a relatively good state of preservation for australopithecine fossils from South African sites. The only other reasonably complete and well-preserved adult *Australopithecus* cranium is that of Sts 5 (‘Mrs. Ples’), which is undistorted but unfortunately has no teeth preserved. The Taung child, type specimen of *A. africanus*, is that of an infant, and although very well preserved and probably complete before it was blasted out by lime mining, it is now missing parts of the braincase and mandible. Other *Australopithecus* crania from Sterkfontein are either partial or very badly crushed, or both. These include the first discovered adult specimen TM 1511, Sts 17, Sts 19, Sts 52, Sts 71, StW 13, StW 53, StW 252, StW 505, and StW 576. Although Clarke (1990) conducted a programme of cleaning and reconstruction of several Sterkfontein and Swartkrans fossils in the Transvaal Museum (now the Ditsong Museum), there remain many requiring such attention. The effects of pressure on crania within cave sediments, as well as shearing effects through collapse of deposits, were described and illustrated by Brain (1981, pp. 134-136). These effects are well seen by comparison of StW 573 with a large *Paranthropus* cranium, SK 83 (Fig. 11). In facial view, both display a similar deformation of the frontal pushed downward into the orbits. But this condition was produced by difference processes. In the case of SK 83, it was pressure of sediments, including rocks, from above that crushed the braincase downward, whereas in StW 573 the whole lower and mid-face was rotated upward into the frontal by a shearing force, most probably shifting or collapse within the talus slope prior to cementation of the deposit.

**Fig. 11.**
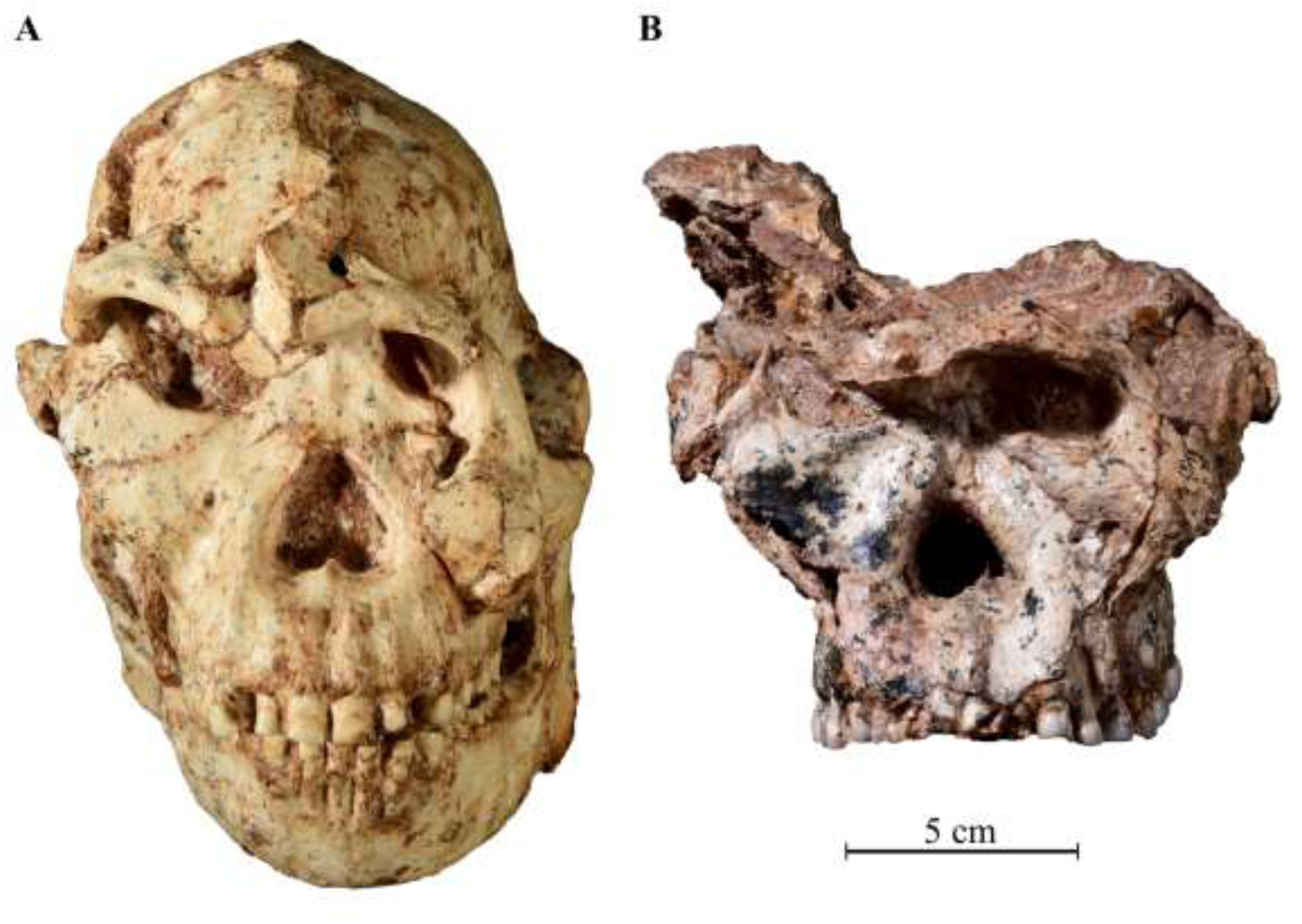
Comparison of differential crushing between StW 573 and *Paranthropus* SK 83 from Swartkrans. The face of StW 573 has been rotated upwards into the frontal from below, while the braincase of SK 83has been crushed downwards from above.

### 4.2. The skull as a whole

Although it is of similar age to earlier fossils of *A. afarensis*, the most comprehensive skull data available for that species is based on the ca 3 Ma A.L. 444-2 from Hadar, Ethiopia (Kimbel, Rak and Johanson, 2004). Like StW 573, that skull consists of the rare association of cranium and mandible in one adult individual. The two skulls have some similarity in overall shape, including depth and robusticity of the zygomatic, temporal lines that sweep strongly medially behind the browridge and meet as a crest on the parietals, bilateral convexity of the nasoalveolar clivus, and possession of a large diastema. They also exhibit some differences. A.L. 444-2 has a different upper facial structure, with relatively narrow infraorbital region and a ‘squared-off’ superolateral corner to the anteriorly prominent supraorbital bar. In A.L. 444-2 in contrast to StW 573, there is a shelf extending over porion to meet with the supramastoid crest. There is no such shelf in StW 573. The mastoid process of A.L. 444-2is large and, in posterior view, projects well below the level of the occipitomastoid crests. This is the reverse of the situation in StW 573. Also, in posterior view, the sides of the braincase in A.L. 444-2 and in A.L. 333-45 slope markedly superomedially, contrasting with the almost vertical sides in StW 573 (Fig. 12). The distance from the posterior margin of the foramen magnum to the external occipital protuberance is short in A.L. 444-2and considerably longer in StW 573.

**Fig. 12.**
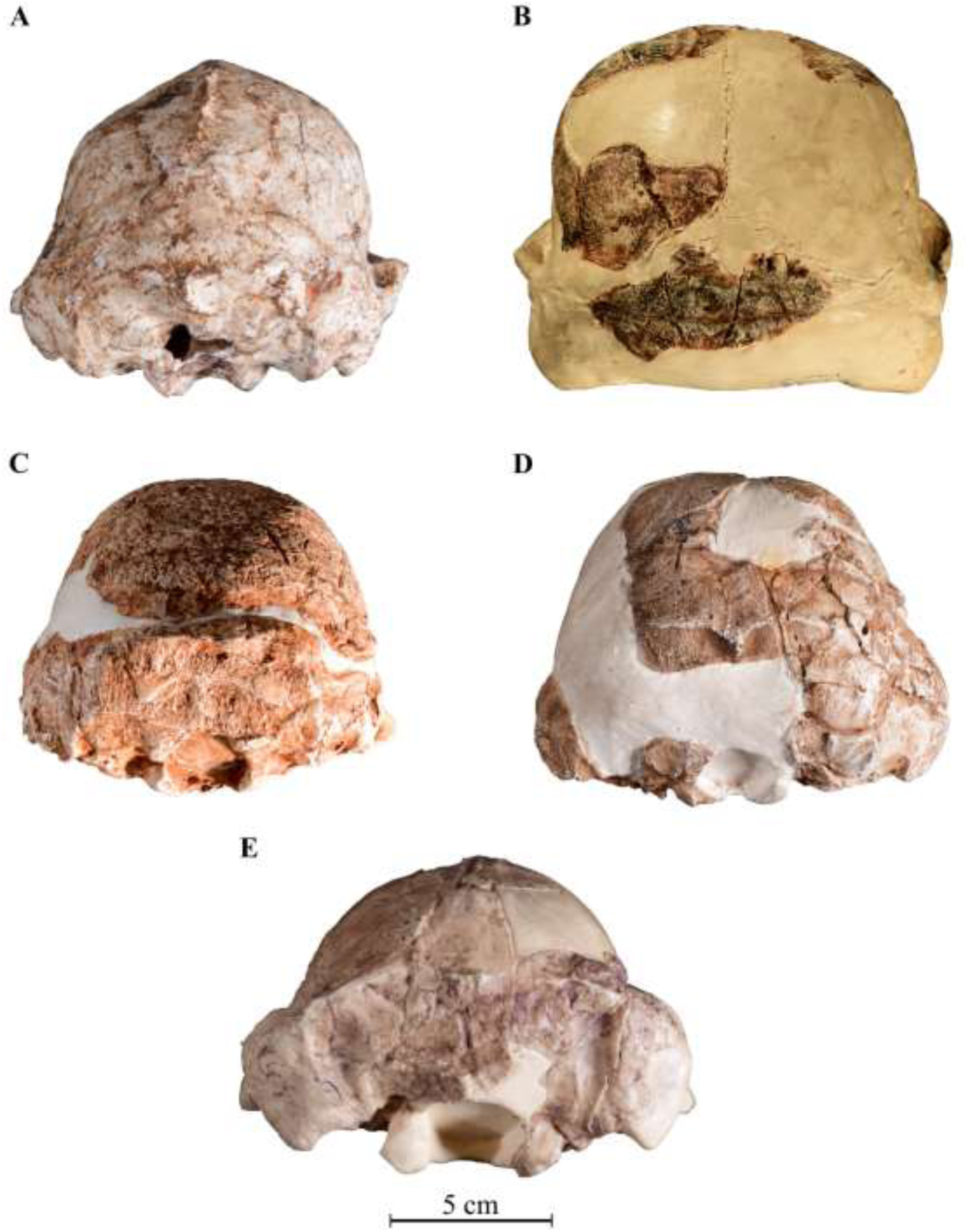
Posterior view of braincases. A) StW 573; B) StW 252; C) Sts 5; D) StW 53; E) A.L. 333-45. Note that in StW 573 and StW 252, the sides of the braincase are more vertical and curve around parietal bosses. The sloped appearance of the left side of StW 573 is due to warping.

### 4.3. The face

The form of the frontal squame and supraorbital region of StW 573 closely matches that of the Sterkfontein *Australopithecus* crania StW 252 and StW 505, exhibiting a low, slightly concave anterior squame above glabella and temporal lines that sweep medially behind the brow ridge. In this they contrast strongly with other Sterkfontein crania, StW 53 and Sts 5, which have higher convex frontal squama with a metopic ridge and temporal lines that do not sweep so sharply medially behind the brow ridge.

Lockwood and Tobias (1999) described the glabella region of StW 505 as differing from Sts 5 and StW 13 in not having a midline peak but instead being composed of twin superciliary eminences divided by a midline depression. They stated that such a midline depression was present in TM 1511 and in StW 252, but that the glabella mound is less prominent in those two fossils. The morphology of that region in StW 252 is indeed strikingly similar to that of StW 505, with twin superciliary eminences but not so pronounced because it is juvenile. As far as TM 1511 is concerned, the surface of the region is damaged and partly missing, but there is enough preserved to show that it did not have the same morphology with twin superciliary eminences as seen in StW 505 and StW 252. That region is missing also in StW 53. The form of the supraglabella surface in StW 573, even though cracked and slightly displaced, appears to have had a browridge of even thickness with the superciliary portion curving down to glabella on each side but separated from each other by a ca 15 mm broad concave area. As the bone on either side of glabella is crushed, it is difficult to determine whether there were superciliary prominences.

StW 573 has a great interorbital breadth (24 mm from dacryon to dacryon; Table 2), which approaches the 27 mm given by Lockwood and Tobias (1999, p. 645) for StW 505 for the breadth between the medial orbital margins. StW 252 also has a great interorbital breadth of at least 20.4 mm. There is a flat midface on either side of the inferior end of the nasal bones. This morphology is similar to that of *Paranthropus* and in marked contrast to that shown by the Sterkfontein specimens Sts 5, Sts 17, Sts 52, and StW 53 (Fig. 13), as well as the Taung child. They all display a narrow interobital distance (ca 16.0 mm in Sts 5 and an estimated 15.5 in StW 53) and an anteriorly prominent midface.

**Fig. 13.**
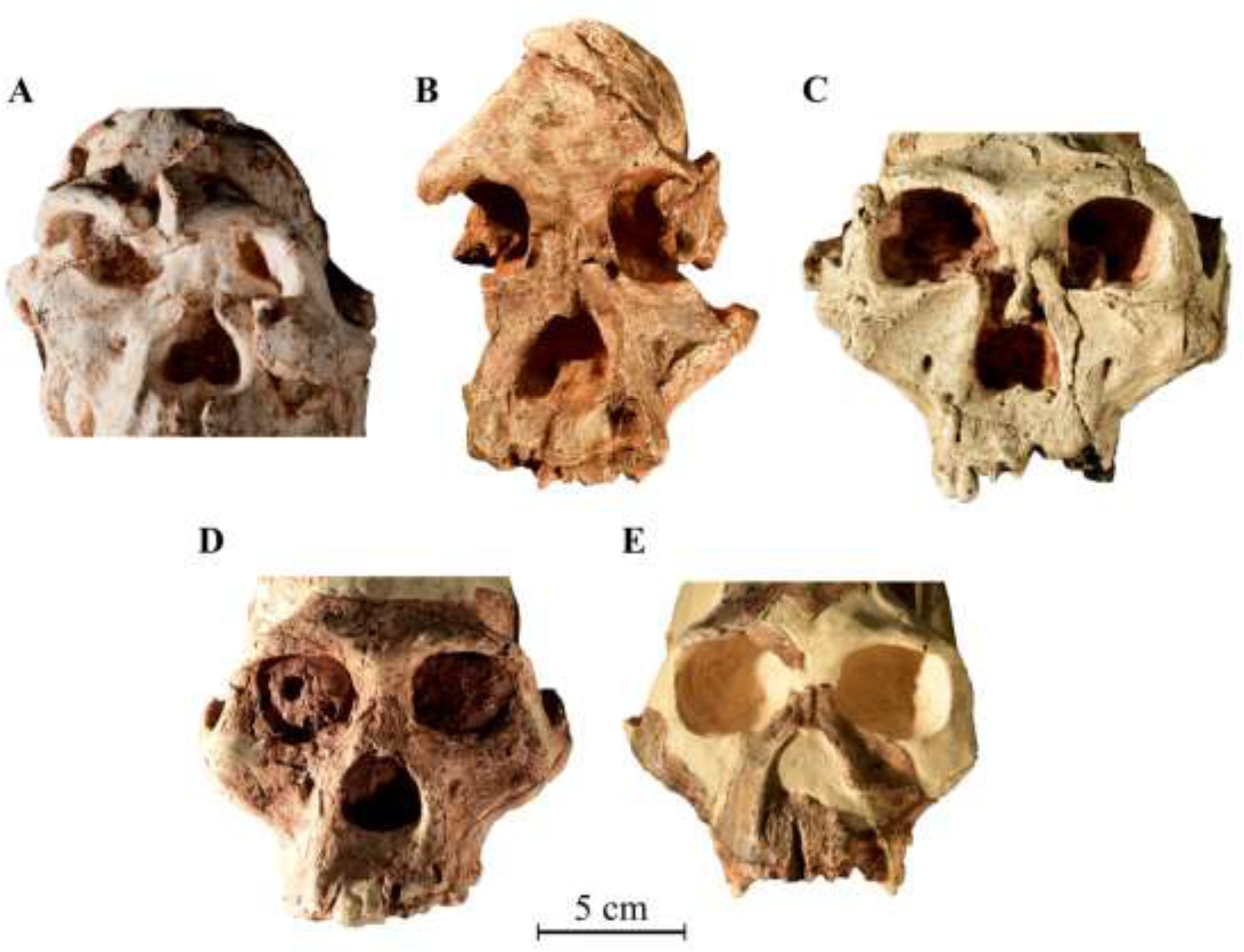
Comparison of interorbital distance in: A) StW 573; B) StW 505; C) SK 48; D) Sts 5; and E) StW 53.

The lower nasal margin and nasoalveolar clivus morphology of StW 573 (Fig. 14) aligns it with some Makapansgat and Sterkfontein fossils, namely MLD 9, StW 183, and StW 498, as well as with *Paranthropus* (e.g., SK 12). They all show a gently concave prenasal fossa passing smoothly into the floor of the nose, and in those *Australopithecus* specimens the clivus is bilaterally convex. Other fossils such as TM 1512, Sts 5, StW 53, Sts 17 and Sts 53 have more sharply defined inferior nasal margins. Some have a flat or bilaterally concave clivus set between the pronounced canine pillars. Others have a convex clivus, e.g., TM 1512 and Sts 52. Kimbel, Rak and Johanson (2004) mentioned this variability in the Sterkfontein sample by commenting that the surface of the clivus is frequently flat but sometimes sunken between the canine pillars. They noted a contrast between Sts 5 and StW 73 as compared to TM 1512 and Sts 52a. Those latter two specimens have a slight convexity to the clivus. In StW 573, there is a bilateral convexity of the clivus, which projects anterior to the canine pillars and is emphasized by the pronounced ridges of the incisor root sockets. It does not conform with the less convex clivus exhibited by TM 1512 and Sts 52a. In those fossils, the incisor sockets are more vertically emplaced and are not very pronounced. Also they do not have an incisor-canine diastema, and the clivus does not pass so smoothly into the nasal floor but is marked by a slight sill. StW 573 has very procumbent incisors and a wide diastema. In this region it contrasts also with a newly reconstructed maxilla with complete dentition (StW 576) that has vertically emplaced incisors, no incisor root prominences, and no diastema (Fig. 15).

**Fig. 14.**
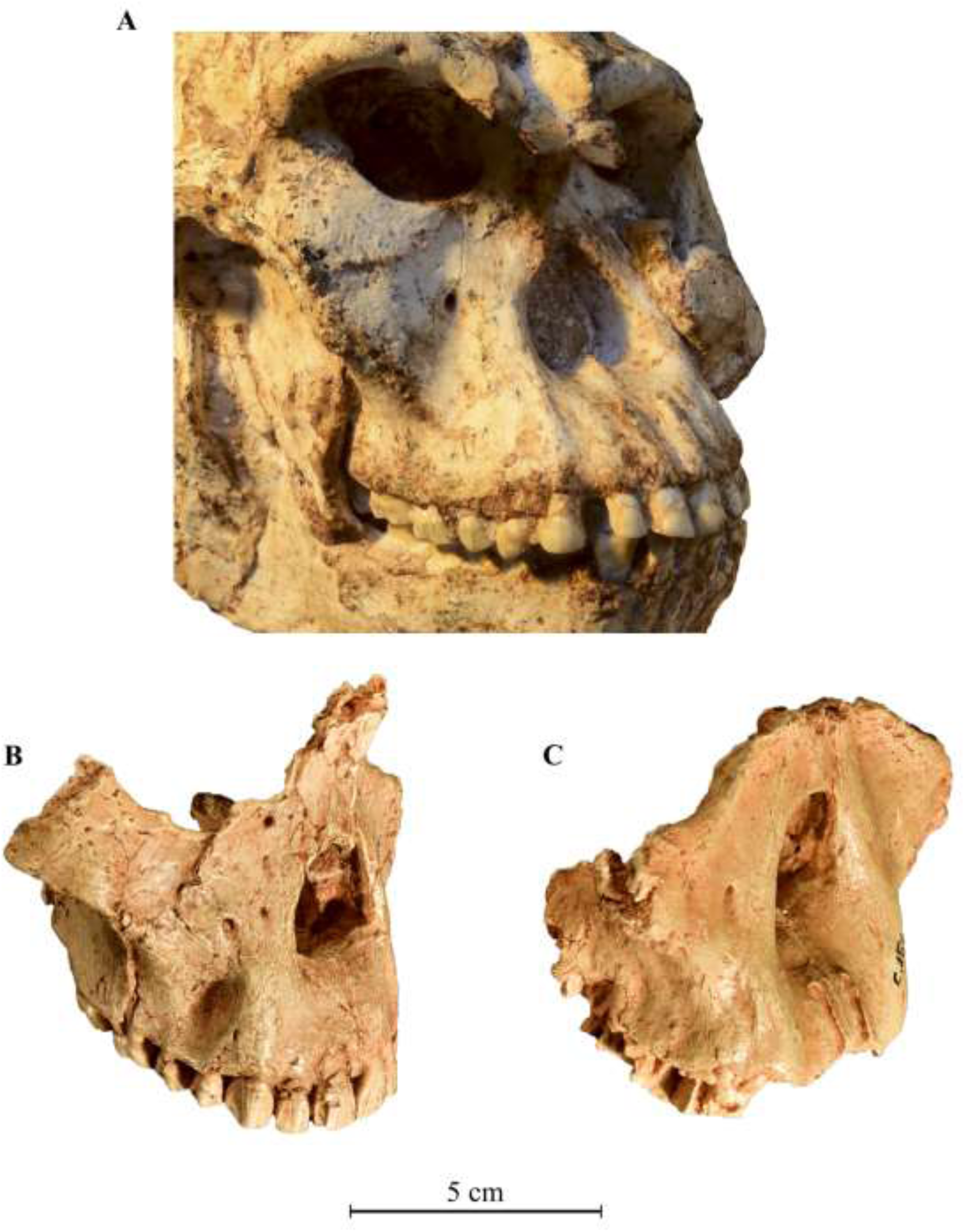
Three-quarter views of the midface of: A) StW 573; B) Sts 52a; and C) SK 12. It shows the concave lower nasal aperture in StW 573 and SK 12 (*Paranthropus*), and the sharp lower nasal margin in Sts 52 (*A. africanus*). This figure also shows the presence of a large diastema in StW 573 and lack of a diastema in Sts 52.

**Fig. 15.**
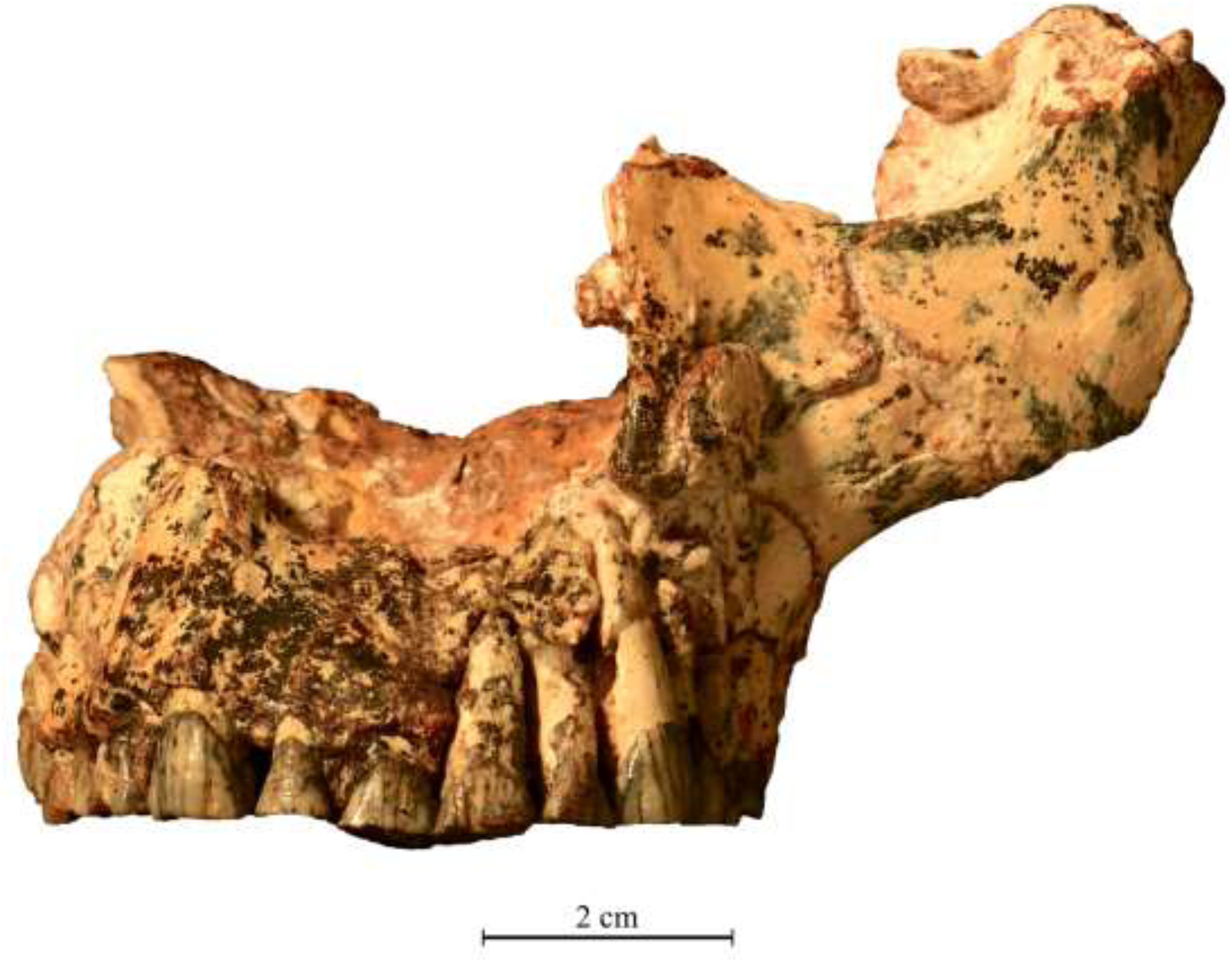
Facial view of the StW 576 *A. africanus* maxilla. It shows the sharp angulation of the inferior margin of the zygomatic process of the maxilla and the lack of a diastema between canines and lateral incisors.

The clivus of StW 505 is damaged laterally by crushing and by preparation gouge marks on the inferior nasal border. Thus it is difficult to make comparisons. The observation by Lockwood and Tobias (1999) that there are strong ridges running diagonally from the inferolateral nasal aperture to the ‘conjoint central and lateral incisor juga’ needs to be considered together with the damage. It seems possible that on each side these ridges were originally marking the surface of the lateral incisor sockets but have been emphasized and given a somewhat diagonal appearance as a result of the crushing and deformation on each side. Prior to the crushing, this region could well have resembled that of StW 573 in the form of a bilaterally convex clivus. It certainly has the same deep midsagittal groove between the central incisor sockets. Such morphology does not resemble that of the other Member 4 specimens listed by Lockwood and Tobias (1999) as having the same diagonal ridges, i.e., TM 1512, Sts 52, Sts 53, StW 73, and StW 391. Those specimens conform with the morphology of Sts 5, StW 53, and StW 576. They do not have marked diagonal ridges or crests. If the StW 505 diagonal ridges determined by Lockwood and Tobias are indeed not emphasized or oriented by the crushing, then their association with the central incisors is probably related to the importance of those incisors in food preparation, as will be discussed below for the dentition of StW 573.

The inferior margin of the StW 573 zygomatic process of the maxilla takes off above the first and second molars and rises more superiorly than laterally to meet the base of the well-defined masseteric attachment. In this it strongly resembles StW 505 and contrasts with the more laterally directed inferior margin of the zygomatic process exhibited by StW 53, Sts 5, and StW 576 (Fig. 11). The difference in the angulation of the inferior margin of the zygomatic is probably related to the much longer faces of StW 505 and StW 573, as well as to the greater robusticity of the zygomatics in those specimens.

The large diastema seen in facial view of StW 573 is characteristic of the great apes and *A. afarensis* (White et al., 1981) and is seen also in the Sterkfontein specimens StW 252 and StW 183. It does not occur in TM 1511, TM 1512, Sts 5, Sts 52, Sts 53, StW 73, StW 53, or StW 576. In the apes, the diastema accommodates the long lower canine crown which shears against the anterior face of the upper canine when the jaws are closed. In StW 573 the lower canines do not lodge in the diastema, but instead they occlude against the upper canines. The need for the diastema is only apparent in the infant dentition, a good example of which is the TM 1516 symphysial fragment with unerupted canine (Fig. 16). If that unerupted canine is placed in what would be its occluded position superimposed on the StW 573 mandible, it can be seen that it fills the diastema. Hence the diastema is a retained ancestral condition which is necessary for the large canine of the young but becomes unnecessary in the adult when the canines are worn down on their tips.

**Fig. 16.**
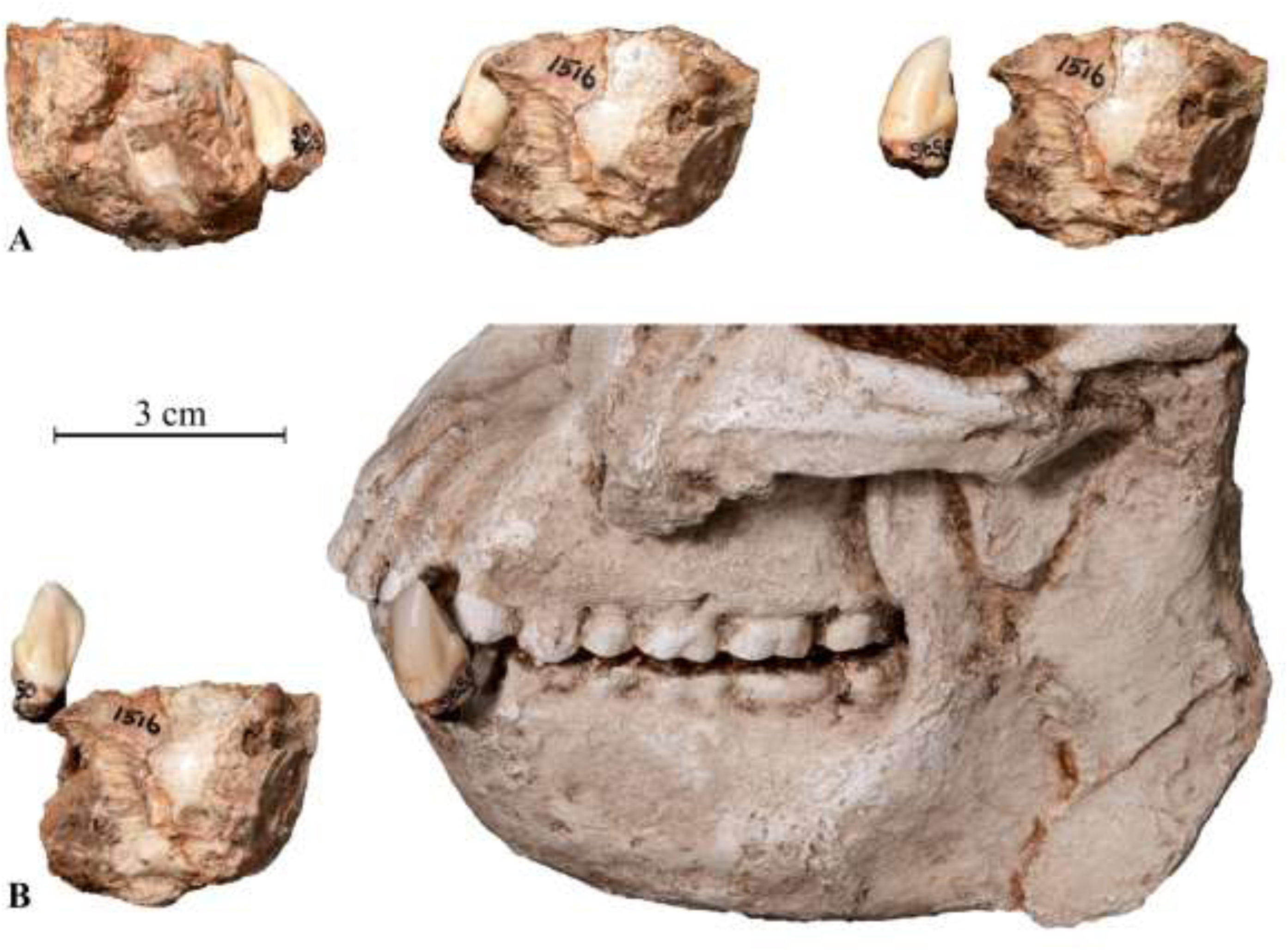
TM 1516 infant mandible, showing: A) the unerupted canine in its crypt and separated; B) the lower canine placed in erupted position relative to its mandible and superimposed on StW 573 mandible to occupy the diastema.

### 4.4. The braincase

The profile of the StW 573 braincase in posterior view, with its almost vertical sides that turn superomedially around parietal bosses, is very different from the profiles of *A. afarensis* (e.g., A.L.- 442) and the *A. africanus* male StW 53 in which the sides slope superomedially from the supramastoid crest. Instead, StW 573 resembles MLD 1, the type specimen of *A. prometheus*, as well as the juvenile StW 252, which also belongs to *A. prometheus* (Clarke, 2013). In those specimens, the sides of the braincase are almost vertical and turn superomedially around parietal bosses. The parietal profile is only subtly distinguishable from that of Sts 5, but StW 573 does differ in the more inferior orientation of its nuchal plane, in the inferiorly prominent occipitomastoid crests, and in its possession of a sagittal crest. In that latter feature, StW 573 aligns with StW 505. The uncorrected, and therefore minimum, endocranial volume of StW 573 is 408 cc (Beaudet et al., in press). This is clearly considerably smaller than that of StW 505, which Holloway (2012) has estimated at slightly above 575 cc. Holloway (2000) also estimated the cranial capacity of MLD 1, the *A. prometheus* type specimen, at 500 to 520 cc. This big difference in brain size is probably due to the much earlier age of StW 573, as well as its being a female, whereas StW 505 is a very large male.

In lateral aspect, a notable distinction between StW 573 and crania of *A. africanus* is in the size of the zygomatic arch, which is deep in StW 573 18.5 mm anteriorly and 13.0 mm midway along the length (Table 2). In the male *A. africanus* StW 53, those dimensions are 11.5 mm and 8.5 mm respectively, and in the female *A. africanus* Sts 5 the dimensions are 12.5 and 8.5 mm respectively. In StW 573 the arch terminates posteriorly by merging into the temporal squame anterior to porion, as it does also in StW 505. That region differs in Sts 5, StW 53 and Sts 19, where the arch continues posteriorly as a shelf over porion until it merges with the supramastoid crest.

### 4.5. The palate and basicranium

The palate of StW 573 is long, narrow and shallow (Table 3), with a shallow slope from the incisive foramen to the incisor alveoli. The palate tapers anteriorly such that the breadth measurement at the level of P^3^ is about 30 mm. Kimbel et al. (2004) observed that *A. afarensis* has the narrowest, shallowest palate of any *Australopithecus* species, except perhaps *A. anamensis*. The measurements of StW 573 in Table 3 show that it is close to *A. afarensis* in palate length and depth, in which it differs from the other three Sterkfontein *Australopithecus* that are listed. The long and shallow palate of StW 573 is similar to that of StW 252, which also differs radically in shape and in cheek teeth size and morphology from many other Sterkfontein palates (Fig. 17).

**Table 3.** Palate dimensions (in mm) and shape index. Length and breadth measurements of StW 573 [in brackets) are estimated for now as they are partially obscured in the MicroCT scans. Other comparative measurements are taken from Kimbel, Rak and Johanson (2004), Table 5.10.

| **Specimen** | **Palate Length** orale-staphylion | **Palate Breadth** endomolare-endomolare | **Palate Depth** at M1-M2 | **Palate Shape Index** breadth/length x 100 |
| --- | --- | --- | --- | --- |
| StW 573 | [69] | [35] | 11.3 | 50.7 |
| *Sterkfontein Australopithecus* |  |  |  |  |
| Sts 5 | 65.3 | 35.7 | 18.0 | 54.7 |
| Sts 53 | 54.0 | 32.0 | n/a | 59.3 |
| StW 73 | 58.0 | 30.0 | 14.5 | 51.7 |
| Olduvai OH 24 | 50.7 | 35.5 | 15.0 | 70.0 |
| KNM ER 1813 | 53.5 | 32.7 | 16.0 | 61.1 |
| *Paranthropus* |  |  |  |  |
| OH 5 | 79.1 | 38.2 | 21.0 | 48.3 |
| SKW 11 | 60.0 | 34.6 | 15.0 | 57.7 |
| *Australopithecus afarensis* |  |  |  |  |
| A.L. 444-2 | 75.0 | 41.0 | 12.0 | 54.7 |
| A.L. 200-1a | 65.0 | 33.5 | 8.5 | 51.5 |
| *Early Homo* |  |  |  |  |
| A.L. 666-1 | 62.5 | 39.3 | 16.5 | 62.9 |

**Fig. 17.**
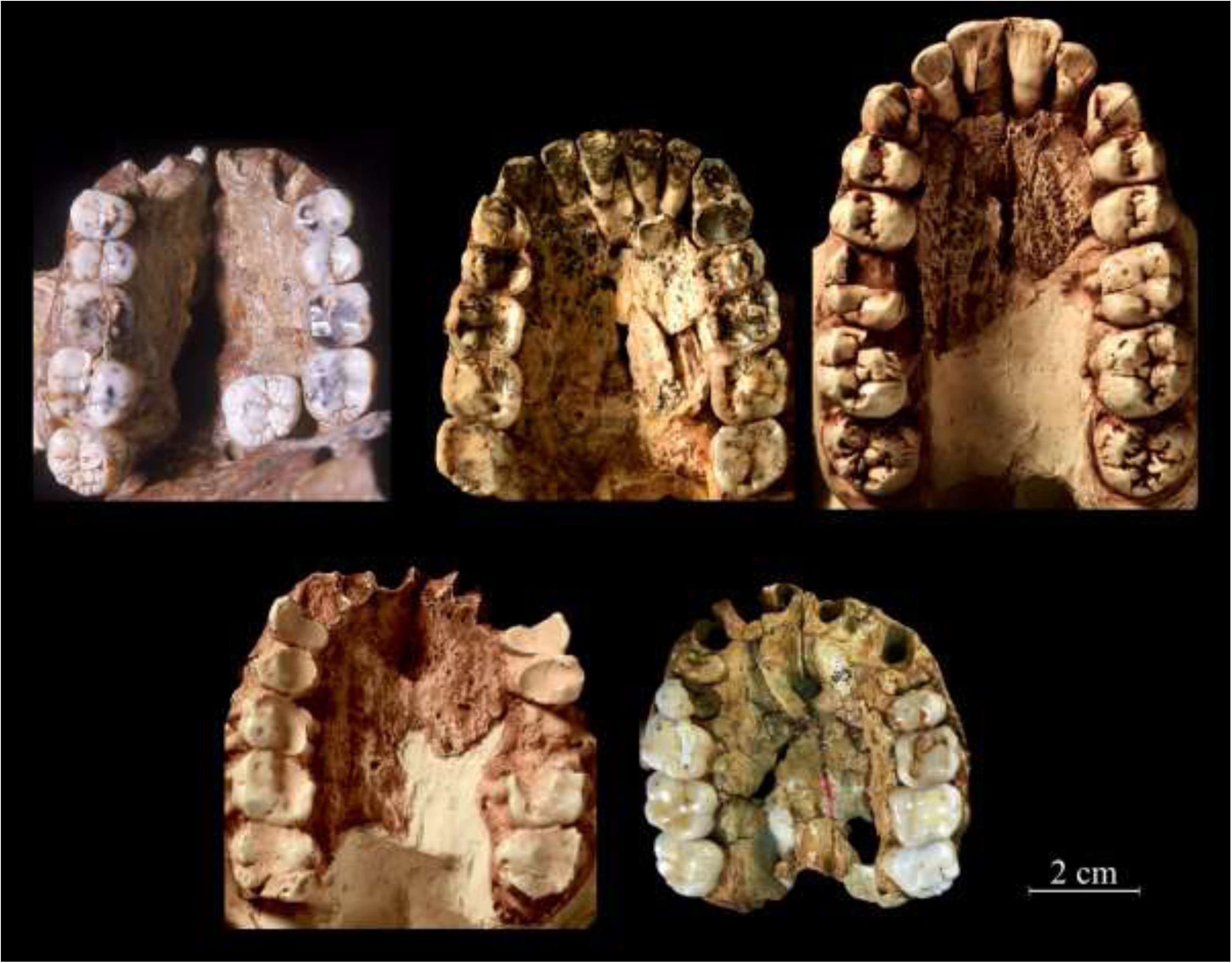
Comparison of *Australopithecus* palates from Sterkfontein Member 4. Top left: TM 1511 *A. africanus*; top middle: StW 576 *A. africanus*; top right: StW 252 *A. prometheus*; bottom left: StW 53 *A. africanus*; bottom right: Sts 53 *A. africanus*. Photos by M. Lotter and R.J. Clarke.

StW 573 has an anteroposteriorly long nuchal plane measuring 48.5 mm from the posterior margin of the foramen magnum up to inion. This nuchal plane is on the same plane as the foramen magnum (see Fig. 2). Sts 5 and StW 53, however, have short nuchal planes (33 mm and 37 mm estimate respectively) that angle superiorly from the plane of the foramen magnum. The occipitomastoid crests of StW 573 are very pronounced and are bordered laterally by deep and wide digastric grooves that run smoothly on to the inferior surface of the tympanic plate. Sts 5 and StW 53 have weakly expressed occipitomastoid crests with modest digastric grooves, anterior to which are well-defined petrous crests forming the base of the tympanic plates, with a vaginal process of the styloid against its posterior face. In their discussion of this region in StW 505, Lockwood and Tobias (1999) noted that a distinct petrous crest was usual in *A. africanus*, but that it did not occur in *A. afarensis*. They considered StW 505 to show the *A. africanus* morphology but noted that it does not have a vaginal process. It can now be seen that StW 573 is closely comparable with *A. afarensis* in this region, having no crest and no vaginal process. Furthermore, it differs from Sts 5 and StW 53 which have crests and vaginal processes (Fig. 18).

**Fig. 18.**
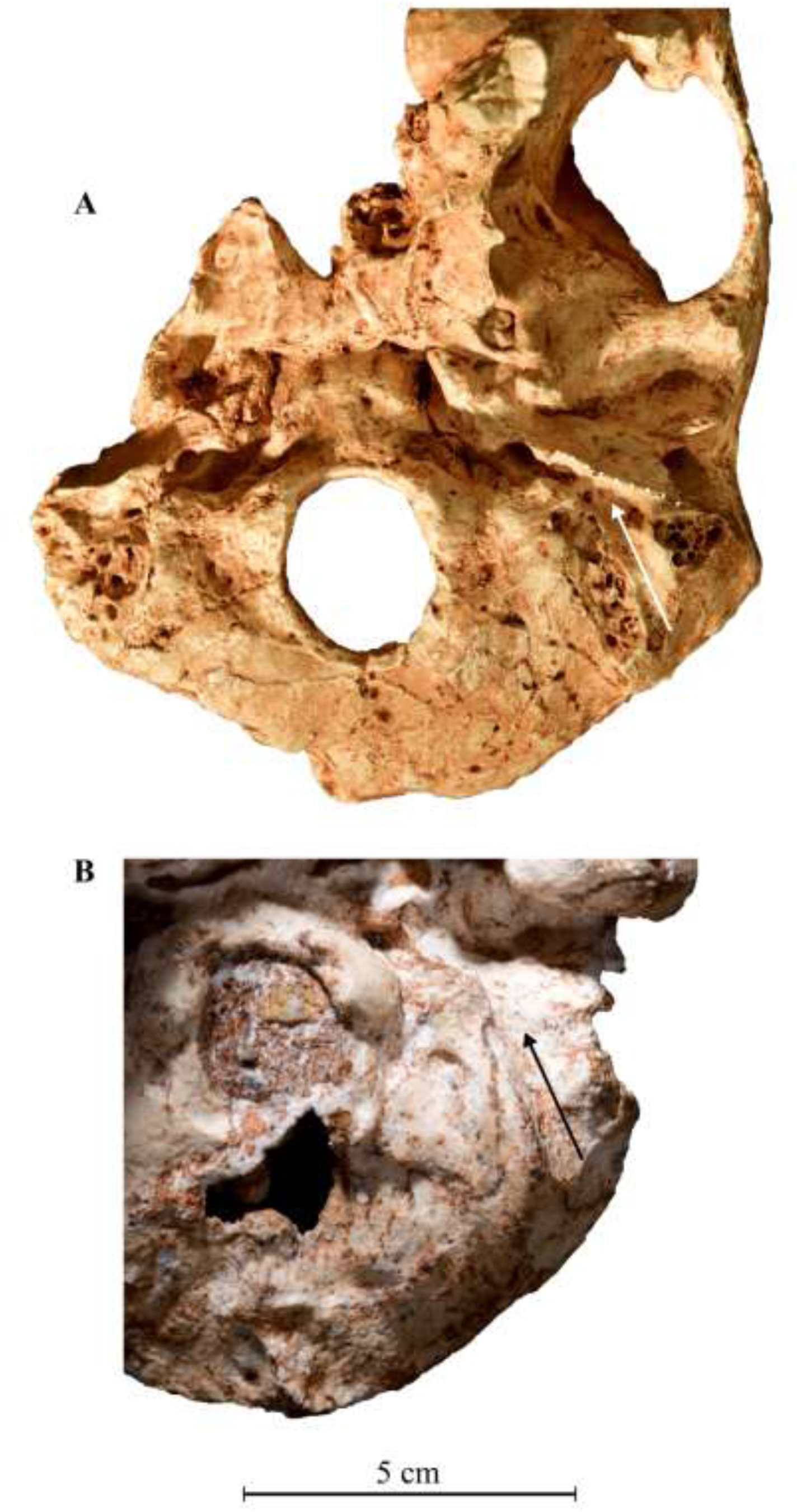
Comparison of the cranial base of Sts 19 (A) with StW 573 (B). It shows the digastric groove (arrowed in A) meeting the sharp crest of the tympanic plate in Sts 19 but merging into a flat tympanic plate in StW 573.

### 4.6. The mandible

The size and morphology of the mandible is extremely similar to the large Sterkfontein Member 4 mandible Sts 36 (see Fig. 19) (Broom and Robinson, 1949), reconstructed by Clarke (1990), but surprisingly, as seen in the scans and measurements, the molar teeth are considerably smaller than those of Sts 36 (see dentition section below). Sts 36 also has heavy wear on the dentition, particularly anteriorly, with P_4_ and M_1_ worn down almost to the alveolar margin. Another very large M4 mandible with extremely heavy wear especially anteriorly is Sts 7 (Fig. 20), which could well have belonged either to the StW 505 cranium or to a very similar large cranium. These mandibles differ noticeably from Sterkfontein and Makapansgat mandibles, such as Sts 52, MLD 18, and MLD 40, in the thickness of their corpus base. Whilst that of StW 573, Sts 36, Sts 7, and StW 498 is relatively thin compared to the upper part of the corpus, the mandibular base of the others is inflated. Thus in StW 573, the anterior part of the base measures about 12 mm, that of Sts 36 is 11 mm, and StW 498 is 11 mm. That dimension in MLD 18 is 16 mm and 19.5 mm in MLD 40. The basal thickness from below M_3_ as far as the gonial angle is 6 mm in both StW 573 and Sts 36, and 2 mm in StW 498; this contrasts with 9.5 mm in MLD 18 and 13 mm in MLD 40.

**Fig. 19.**
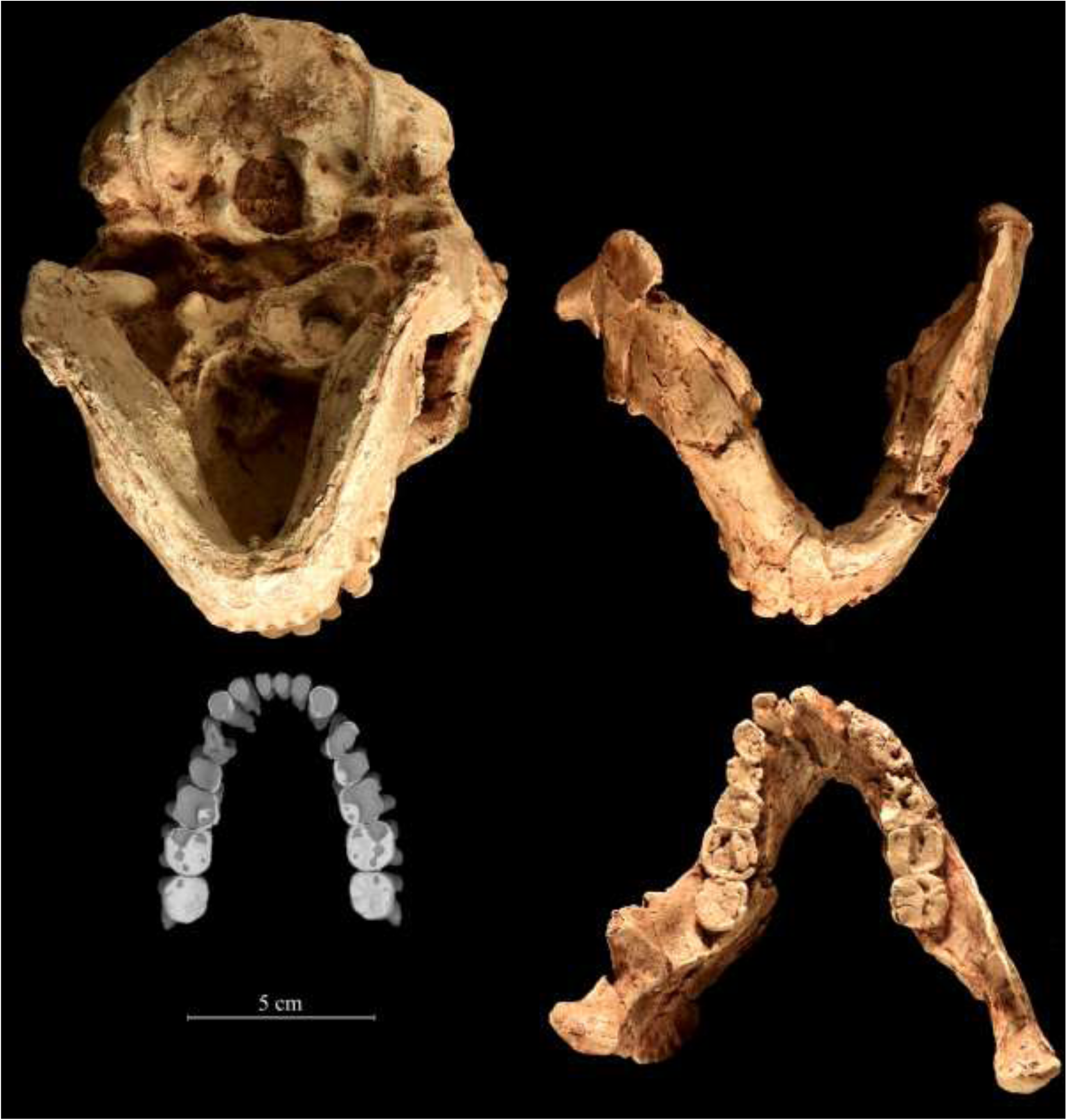
Comparison of mandibular size and shape in A. prometheus from Members 2 and 4. Top left: StW 573 in inferior view. Bottom left: StW 573 scan of mandibular dentition in occlusal view. Top right: Sts 36 in inferior view. Bottom right: Sts 36 in occlusal view. Note the size and shape of the mandibles are the same, but the cheek teeth of StW 573 are very small in comparison to the cheek teeth of Sts 36 from the younger Member 4.

**Fig. 20.**
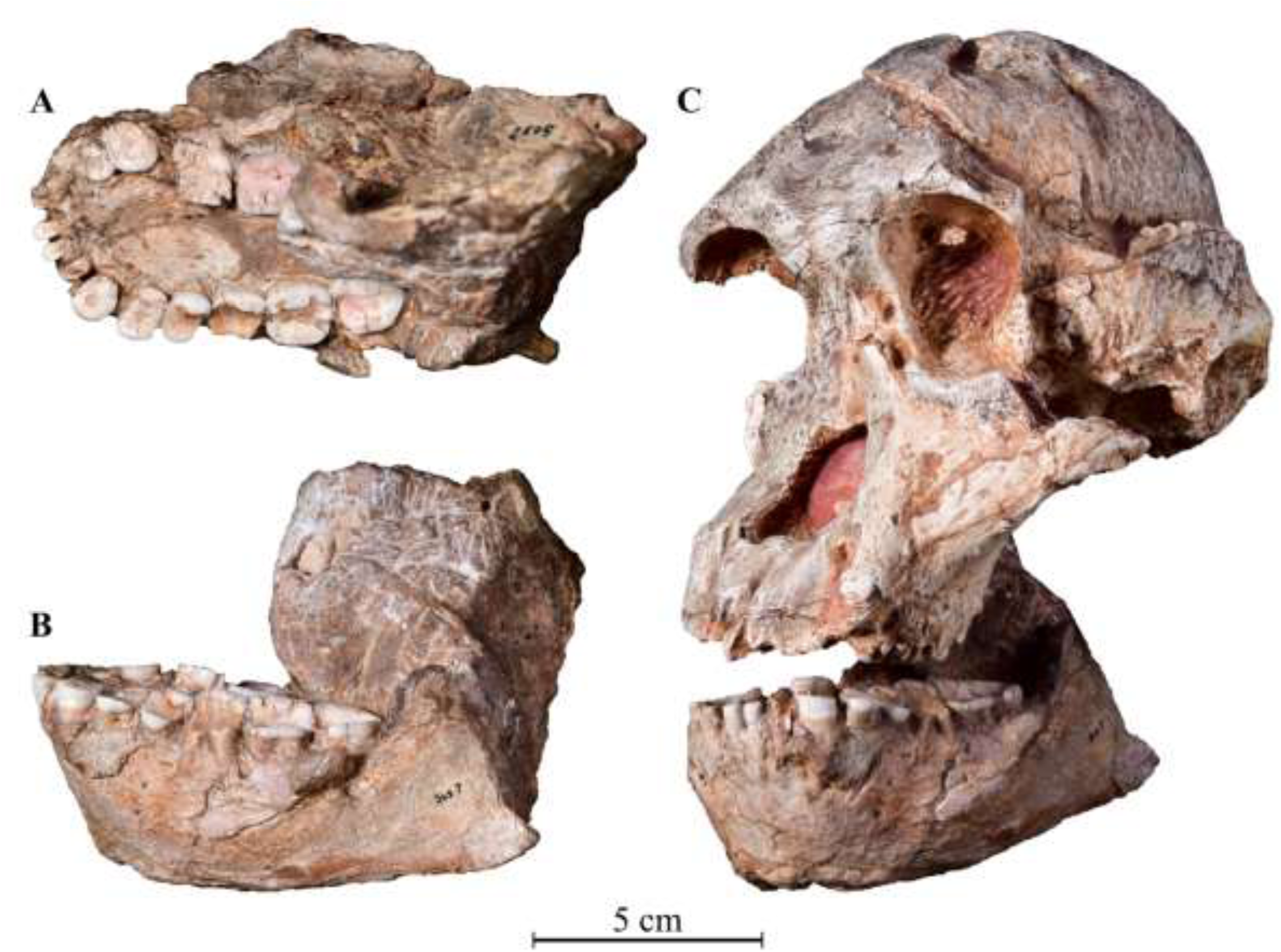
The massive Sts 7 Australopithecus mandible in occlusal view (A), in left lateral view (B), and joined to the StW 505 cranium (C).

### 4.7. The dentition

Although the occlusal surfaces of the StW 573 dentition are not yet fully visible, enough can be seen of the cleaned buccal parts of the upper dentition that, combined with MicroCT scans (Fig. 10) and the exposed lateral and medial views of the dentition to allow for some comparative observations. Three points are of particular note for such comparisons, namely: 1) the large size of the canines and size and shape of their roots; 2) the relatively small size of the cheek teeth set in a large mandible; and 3) the excessive wear on the anterior dentition compared to the moderate wear on the second and third molars.

Prior to the discovery of StW 573, the only other complete upper and lower adult dentition of an *Australopithecus* from South Africa was the Sterkfontein Member 4 specimen Sts 52 maxilla and mandible of a young individual with third molars not fully erupted. The second most complete dentition was StW 252, a full complement of upper teeth of a similar stage of maturity as Sts 52, with third molars not fully erupted. Now, another complete upper dentition, StW 576 from Member 4, has been reconstructed by RJC from a badly crushed partial cranium excavated from grid square R/41 at a depth of about 15 feet below datum on 12 July 1995 (Fig. 17). This specimen is mature, with worn teeth set in a maxilla that extends up to the lower orbit. This fossil proves very important for comparison with the mature dentition of StW 573.

Another palate with complete cheek teeth is the young TM 1511 (Fig. 17) found in 1936 but with two of the teeth having been recovered in recent years from the processing of lime miners’ dumps. The empty sockets for the canines and incisors indicate that they fell out before fossilization. That is the situation also with two other palates that have nearly all of their cheek teeth preserved, i.e., the mature adult StW 53 with heavily worn teeth and the young adult Sts 53 with moderately worn teeth. When these palates are compared in occlusal view (Fig. 17), it is readily apparent that StW 252 differs in many respects from the others, which group together as one morphotype. StW 252 has a much longer palate with large procumbent incisors, a wide diastema, big canines, and very large cheek teeth with bulbous cusps that are situated toward the crown centre such that the sides of the cheek teeth are well visible in this view. In the young adult TM1511, the palate is noticeably smaller than StW 252, the canine and incisor sockets indicate that those teeth were not large, and there is no diastema. The cusps of the cheek teeth are not bulbous and are situated closer to the crown margins such that the sides of the cheek teeth are not visible in this occlusal view. The complete but mature adult dentition of StW 576 shows the same morphology as TM 1511 and provides the extra information in its full complement of incisors that they are not procumbent, the canines are relatively small, and there is no diastema. The palates and dentitions of StW 53 and Sts 53 conform with the morphology of TM 1511 and StW 576, as well as with Sts 52 (See Fig. 8 in Clarke, 1990). In StW 573, the anterior teeth are so heavily worn that comparisons of cusp position and form cannot be made and at present it is difficult to determine much information about the cusps from the scans.

Another very significant feature of StW 573 is the form and size of the canine roots. The MicroCT scans (Fig. 10) show them to be very robust and long, with a marked groove that runs the length of the broad mesial face of the root, which is of oval cross-section. This root morphology and size is seen also in StW 252 (Fig. 21) and in StW 667 (Fig. 22), an upper left canine with broken crown excavated from Member 4. Its root measures 22 mm in length buccally, is 12.5 mm wide buccolingually, and 8 mm mesiodistally. These roots are in stark contrast to the small size and root morphology another Member 4 *Australopithecus*, StW 576 (Fig. 22). In that fossil, the 20 mm long roots are of more rounded cross-section, with only a faint mesial groove, and are relatively narrow (8.7 mm buccolingually and 6.7 mm mesiodistally), in contrast to the wide oval-sectioned roots of StW 573, StW 252, and StW 667.

**Fig. 21.**
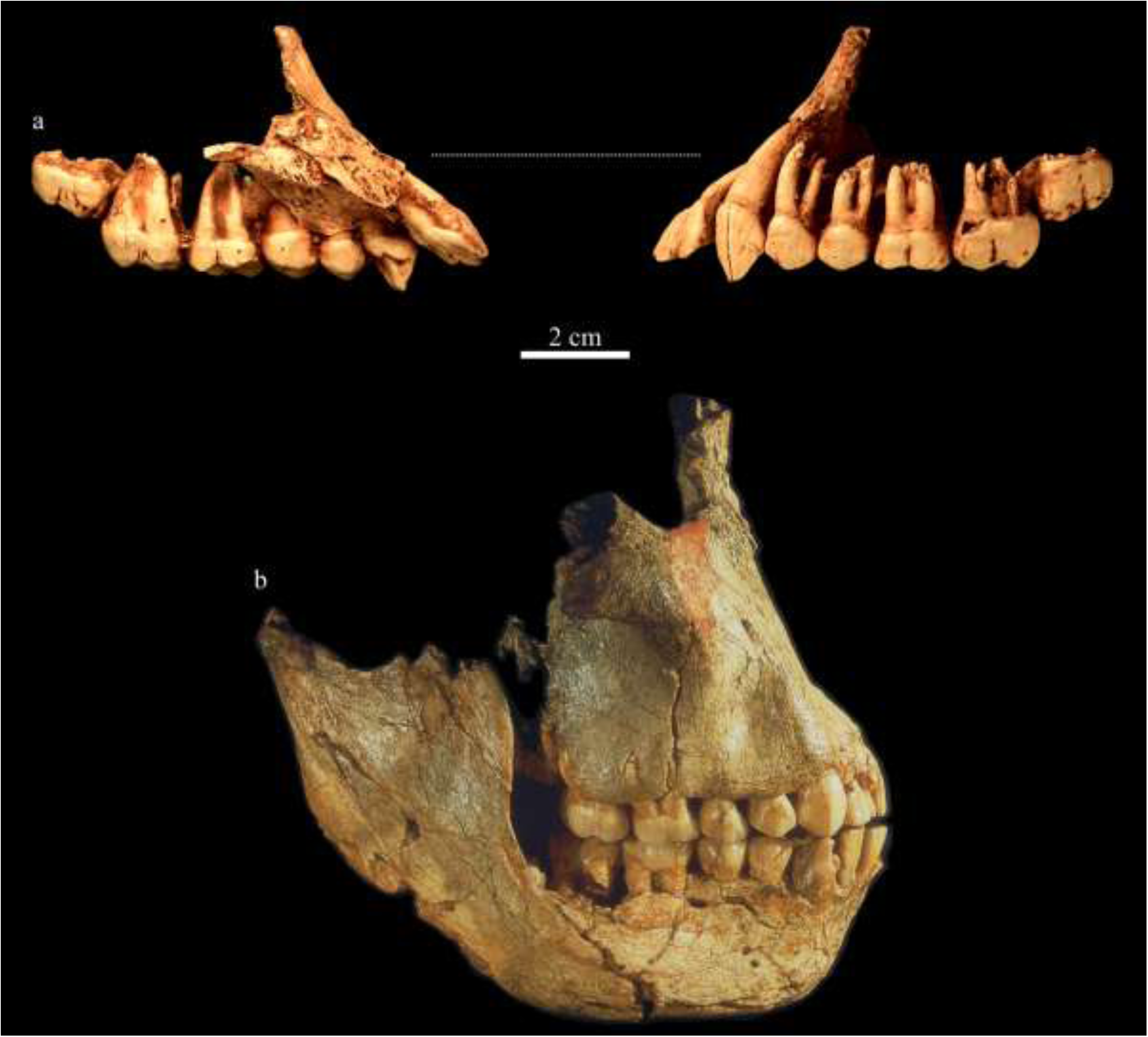
Comparison of incisor procumbence between the two species. a) the lingual view of the left dentition of StW 252, A. prometheus, reconstructed by R.J.C.; b) the buccal view of the left dentition of StW 252, A. prometheus. c) the right lateral view of maxilla with mandible of Sts 52, A. africanus. Note the large canine with massive and broad root in StW 252. Photos by M. Lotter and R.J. Clarke.

**Fig. 22.**
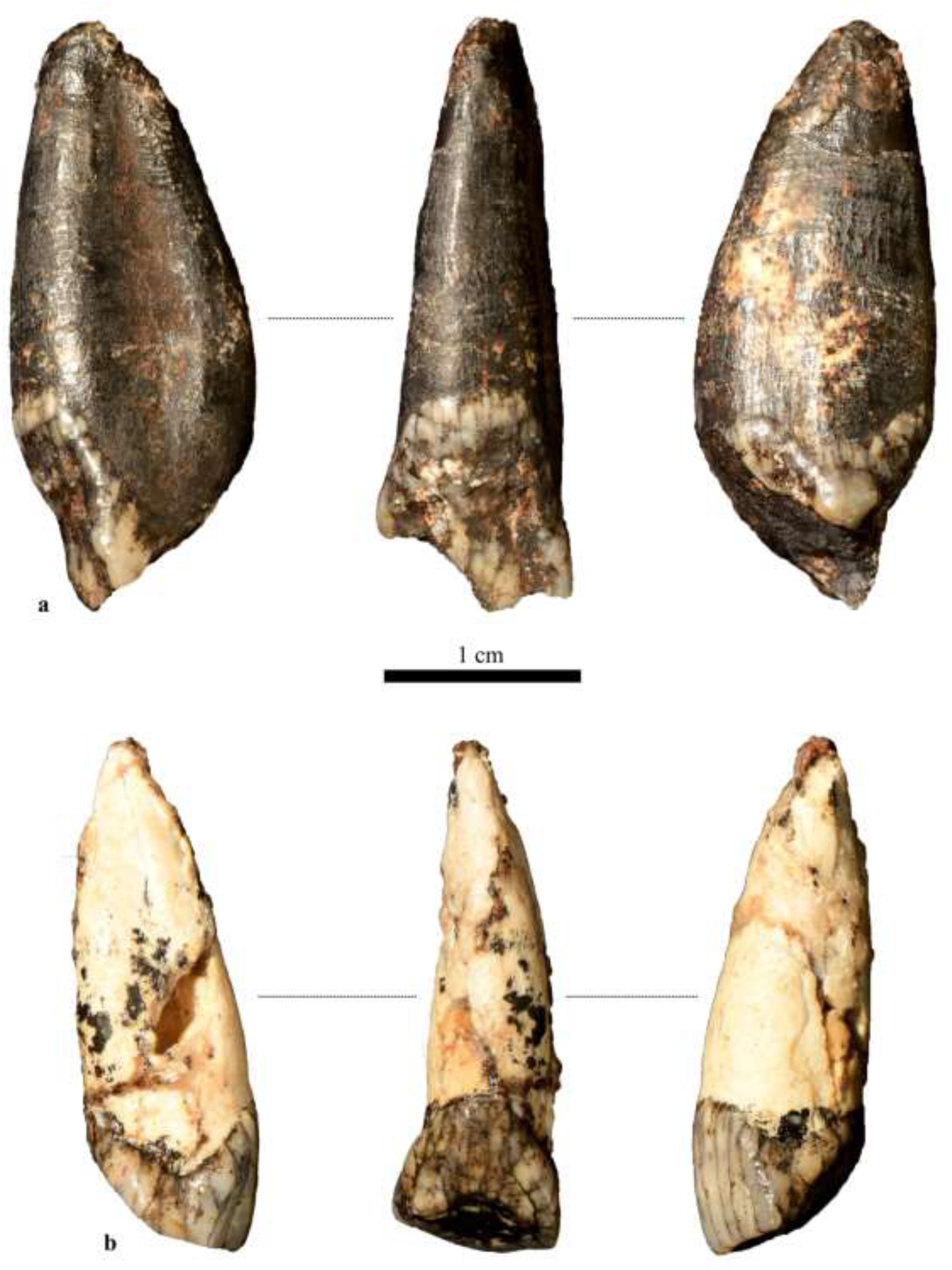
Upper left canines of the two Australopithecus species from Sterkfontein Member 4 compared. a) A. prometheus StW 667, with mesial, lingual and distal views, from left to right. Note that the buccal half of the crown is broken away diagonally. b) A. africanus StW 576, with mesial, lingual and distal views, from left to right. Photos by M. Lotter.

Compared to the large anterior dentition, the cheek teeth of StW 573 are of surprisingly small dimensions (Table 5). The mandible is of the same size and form as that of Sts 36, and yet, as Table 5 shows, the cheek teeth of StW 573 are considerably smaller. Although it is possible that this represents a gender difference with StW 573 being female and Sts 36 being male, that would not explain why the mandibles are of the same size and morphology in both of them. Another possible explanation is that it represents an evolutionary development over time, with the earlier StW 573 having a large mandible with small cheek teeth and the later Sts 36 having retained the large mandible but having also evolved much larger cheek teeth. A second specimen from Member 4 that shows a massive mandible with large, heavily worn teeth is Sts 7 (Broom, Robinson and Schepers, 1950) (Fig. 20).

A third notable feature of the StW 573 dentition is the excessive wear concentrated on the incisors, canines, premolars, and first molars, which form a battery of heavily worn and strongly sloping surfaces consisting largely of dentine (Fig. 10). By contrast, the second and third molars are much less worn. This could be considered as relating to the maturity of the individual as similar heavy sloping wear is seen on the premolars and first molar of the male *A. africanus* cranium StW 53 (Fig. 17). That cranium does not have any incisors and canines preserved for comparison, but the newly reconstructed *A. africanus* maxilla StW 576 (Fig. 17) does have those teeth and shows a different overall pattern of wear to that of StW 573. Although the second and third molars of StW 576 are at an equivalent stage of wear to those of StW 573, the premolars and first molars are well worn to a flat, horizontal surface and the canines and incisors show only moderate horizontal wear. In other words, the canines and incisors of StW 576 do not display the same kind of heavy usage seen in StW 573. The *A. africanus* mandible MLD 18 is at the same stage of wear as StW 576 and also has flat, non-sloping wear on the incisors, canines and premolars (Fig. 23). In this it is in marked contrast to the excessive sloping wear in StW 573.

**Fig. 23.**
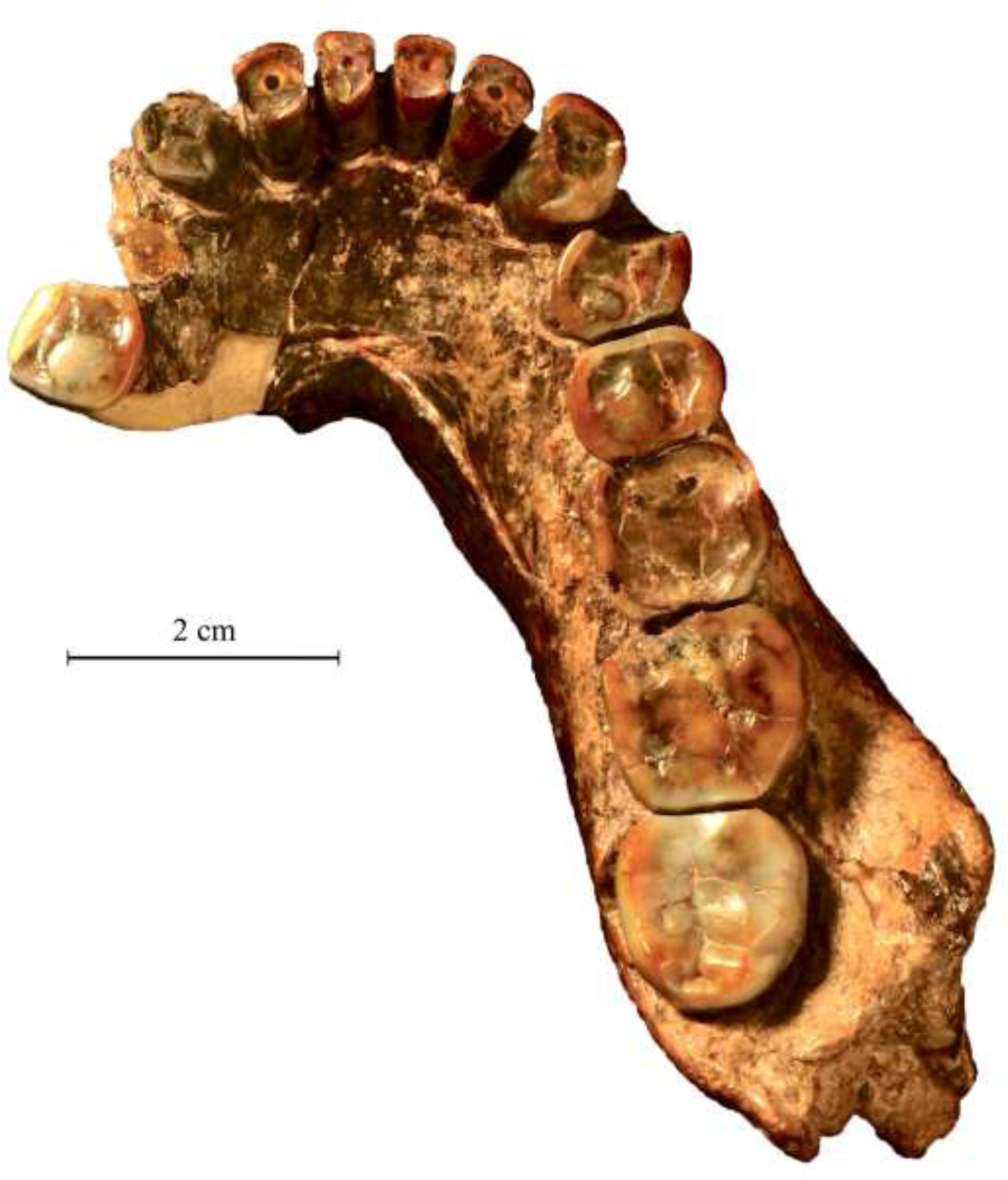
MLD 18 *A. africanus* mandible from Makapansgat in occlusal view. It shows flat occlusal wear on the anterior dentition and more even wear over the whole occlusal plane. Contrast this wear with that of *A. prometheus* in Fig. 10. Photo by M. Lotter.

The array of large and heavily worn anterior teeth in StW 573, contrasting with the much less worn posterior teeth, suggests a specialised feeding function. The same kind of unusual differential tooth wear was recorded for *A. anamensis* (Ward et al., 2010) which those authors found to differ from the tooth wear pattern of the geologically younger *A. afarensis*. They concluded that there was an adaptive shift in diet over time which involved altered use of the anterior dentition in food processing. Such a difference in anterior tooth use is now also seen between *A. prometheus* (StW 573) and *A. africanus* (StW 576) (Fig. 24). In this *A. africanus* specimen, as well as in the MLD 18 mandible, there is no differential excessive anterior tooth wear, but instead a more even wear affecting all teeth, and there is no diastema. These two species do not form an ancestor-descendant lineage but are contemporaries in Sterkfontein Member 4 and Makapansgat, indicating a different feeding behaviour between them. The StW 573 *A. prometheus* wear pattern contrasts also with that seen in *Paranthropus* which has large molars and premolars but with relatively small canines and incisors, no diastema, and no differential emphasis on anterior tooth wear.

**Fig. 24.**
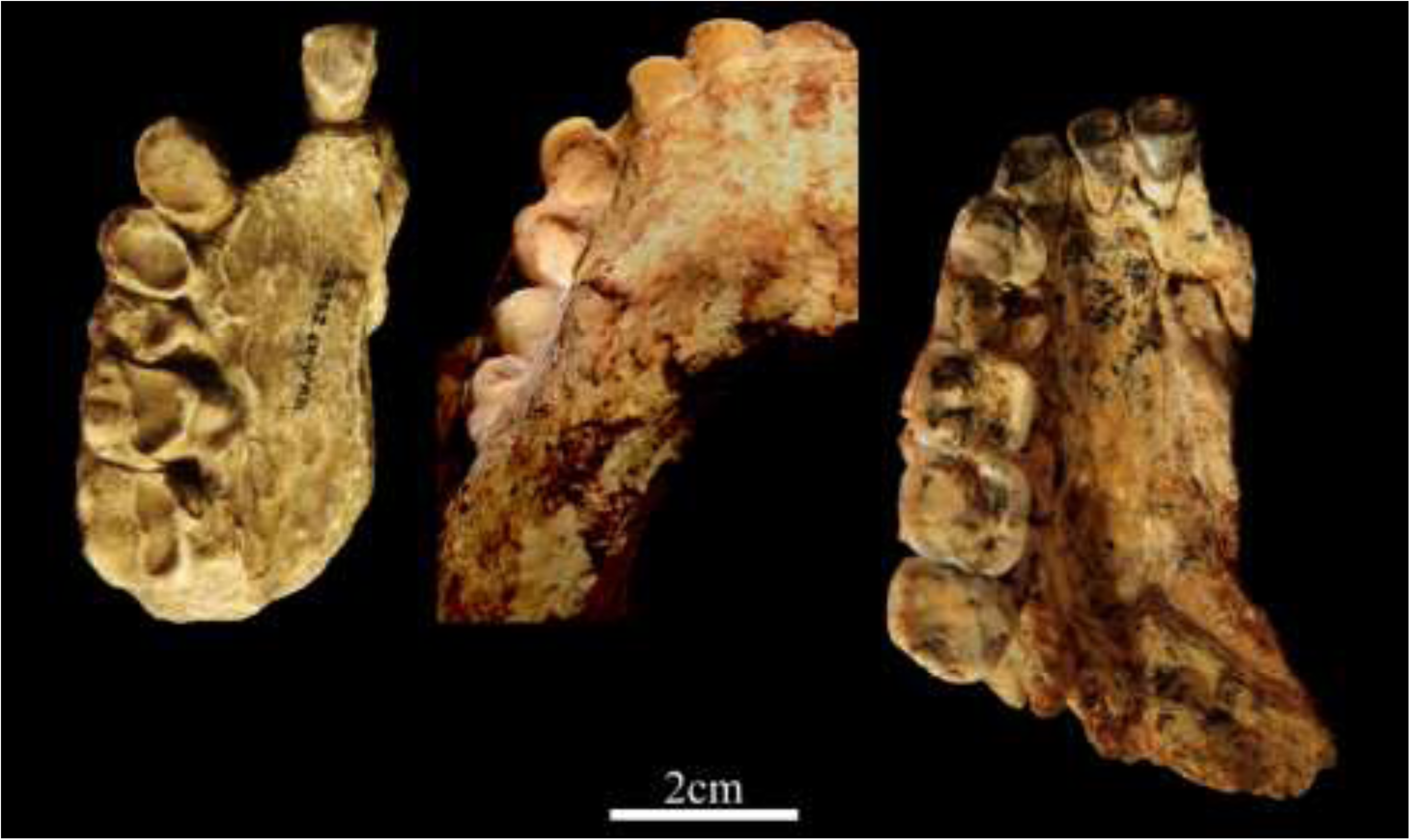
Excessive anterior tooth wear in right upper dentitions of *A. anamensis* and *A. prometheus* contrasted with the more even tooth wear of *A. africanus*. Left: *A. anamensis*, KNM-KP 29283; centre: *A. prometheus*, StW 573; right: *A. africanus*, StW 576. Photos by M. Lotter and K. Kuman.

## 5. Taxonomy

When Dart (1925) published the new genus and species *Australopithecus africanus*, he did not provide a formal definition of that taxon, which would have been limited to the single Taung child skull. It was only 29 years later, following discovery of many adult cranial fossils from Sterkfontein and Makapansgat, that definitions were provided, first by Robinson (1954, 1962) and then by Le Gros Clark (1964) and Tobias (1967). Although Robinson (1954) separated *Paranthropus* from *Australopithecus* at generic level, he considered that the fossils from Taung, Sterkfontein and Makapansgat all belonged not only to *Australopithecus*, but to the species *A. africanus*. Into that species he also placed the *Australopithecus* maxilla from Garusi (Laetoli, Tanzania) that later would be recognised as part of the hypodigm of *Australopithecus afarensis* by Johanson, White and Coppens (1978). Tobias (1980) echoed Robinson by suggesting that the *A. afarensis* fossils were actually East African forms of *A. africanus*. Le Gros Clark (1964) provided a provisional definition of *Australopithecus* based, as he said, on ‘the assumption that all the australopithecine remains so far discovered represent a single genus, *Australopithecus*.’ Not everybody subscribed to that assumption, as it was argued by Robinson (1954) that *Paranthropus* was a genus distinct from *Australopithecus*. Tobias (1967) provided a brief definition of *A. africanus*, but like Le Gros Clark, he included *Paranthropus* in that genus, and like Robinson and Le Gros Clark, he included all the fossils from Taung, Sterkfontein and Makapansgat in the species *A. africanus*. This has long been the generally accepted taxonomy of the South African australopithecines. For example, in an assessment of the phyletic position of *A. africanus*, White, Johanson and Kimbel (1981) wrote: ‘We subscribe to the view that the hypodigm of *A. africanus* includes all known hominid fossils from Taung, Sterkfontein Member 4 and Makapansgat Members 3 and 4.’

A different interpretation was provided by Clarke (1988). Following his reconstruction of the Sterkfontein *Australopithecus* cranium StW 252, he observed that it had notable *Paranthropus*-like differences from *A. africanus* as represented by Taung and Sterkfontein fossils such as Sts 5, Sts 17, and Sts 52. He concluded that StW 252 represented a second species contemporary with *A. africanus*. This second species had a slight concavity of the frontal squame behind glabella, a thin flattened supraorbital margin, a marked medial encroachment of temporal lines just posterior to the supraorbital margin, and a concave facial surface at the level of the frontal processes of the maxilla coupled with anteriorly prominent malars. It has large *Paranthropus*-like cheek teeth but differs from *Paranthropus* in having large incisors and canines, and procumbent incisors accompanied by a wide incisor-canine diastema. Into this second species, Clarke (1988) placed Sts 71, which has a similar frontal morphology, anteriorly prominent cheek bones, large cheek teeth, and high gently curving occipital profile. Other contemporary fossils represented as a group by Sts 5, Sts 17, and Sts 52, have a convex frontal squame, reduced medial encroachment of temporal lines postorbitally, slight anterior prominence of the nasal skeleton, relatively small cheek teeth, thickened supraorbital margin, and more sharply angled occipital profile. In this group, he included MLD 6 from Makapansgat and the Taung child, type specimen of *A. africanus*, and in the second species he included the mandibles MLD 2 from Makapansgat and Sts 36 from Sterkfontein.

This was admittedly a first assessment and two of the features, i.e., thickness of supraorbital margin and gently curving occipital profile, can be variable. The other features, however, showed consistency as an increasing number of fossils was examined or were being newly discovered. Thus Clarke (1994) was able to expand the hypodigm of this second species by adding the child mandibular symphysis with large canine TM 1516, cranium StW 505, mandibles StW 384, StW 14, MLD 27, MLD 29, maxilla MLD 9, and P_3_ StW 401. He noted that the occipitoparietal fossil MLD 1, type specimen of *A. prometheus* (Dart, 1948a) almost certainly belonged to this second species but suggested that, as it is difficult to determine affinity on such a fragment, it was open to debate as to whether *prometheus* could be applied to MLD 2 and other fossils of the second species. Later, Clarke (2013) stated his conviction that *A. prometheus* is indeed a valid name that should be applied to the second species contemporary with *A. africanus* in both Sterkfontein Member 4 and in Makapansgat Member 3.

Most researchers have been reluctant to accept the presence of a second species at these sites and have continued to follow Robinson (1954) in lumping all the *Australopithecus* fossils from Sterkfontein and Makapansgat into the same species as the Taung child, i.e., *A. africanus*. Even though some researchers have conceded that there are specimens that suggest more than one species, they have shied away from committing to such a scenario. For example, Lockwood and Tobias (2002) considered StW 183 to possibly represent something different from *A. africanus* but then demurred because it is immature. They also noted the StW 255 temporal to be similar to the *Paranthropus* cranium WT 17000 and concluded that StW 183 and StW 255 were potentially the best indications of taxonomic variation in Sterkfontein Member 4. This is hardly surprising as the StW 255 temporal is actually part of the same StW 252 cranium that originally inspired Clarke to recognise a second species. StW 255 is now catalogued as StW 252o. StW 183 has the same kind of large cheek teeth with bulbous cusps, large canines and incisors, and flat upper central face seen in StW 252. However, Lockwood and Tobias did not pursue this matter any further. Moggi-Cecchi, Grine and Tobias (2006), following a detailed analysis of Sterkfontein teeth, recognised that there may be a species other than *A. africanus* but stated that ‘such species is difficult to characterise on cranial evidence alone’ and said it was crucial to combine information from study of cranial and dental remains in order to test Lockwood and Tobias’ hypothesis and Clarke’s earlier proposal and ‘eventually to characterise better those specimens which may belong to a taxon other than *A. africanus*.’ In fact, Clarke (1988) did in his original claim for a second species combine study of cranial and dental remains to reach his conclusion.

With reference to studies of occlusal microwear, Grine (2013) observed that ‘the Sterkfontein sample exhibits somewhat greater variability in some parameters than in *Paranthropus robustus* and *A. afarensis*.’ Grine et al. (2013) acknowledged that the morphological variability in the Sterkfontein premolar sample and in the combined Makapansgat and Sterkfontein samples is significantly greater than that of the lowland gorilla, but they concluded that this could be due to change over time. Such an explanation had, however, been refuted 15 years earlier when Clarke (1988) wrote: ‘With regard to change through time, the small and large-toothed morphotypes are found in close proximity both high and low in the talus cone, which suggests that one was not ancestral to the other.’

Despite the above-mentioned indications of morphological variations within the South African *Australopithecus* sample, most researchers have continued to refer to them as all belonging to only one species, *A. africanus*, and have published them as such in comparative tables of measurements and morphological characters and the resulting graphs. This is because they have first accepted that all the fossils belong only to one species, *A. africanus*, and then used the totality of morphological and metrical characters within that united assemblage to supposedly define the species. Such acceptance has effectively blurred the morphological distinctions to be found within the South African fossil sample and is a continuation of the trend started by Broom and Dart of considering all the *Australopithecus* fossils from one site to belong to one species and classing larger-toothed specimens as males and smaller-toothed specimens as females.

The appropriate way to identify a fossil species should be first to identify morphological characters which unite or separate individual specimens, and then to group those specimens according to similarities or differences between clusters of morphological features, whilst allowing for expected variations due to gender, immaturity, or a reasonable amount of individual variation. With this in mind, Table 5 provides a list of morphological characters that cluster the South African *Australopithecus* cranial sample into two distinct groups (A and B). This was achieved by first noting similar or different clusters of features on more complete crania and mandibles, then by noting one or more of those distinguishing features on more fragmentary fossils, and then by grouping them accordingly. It so happens that the cluster of morphological characters shown by Group B aligns this set of fossils with the Taung child, the type specimen of *A. africanus*. The cluster of morphological characters shown by Group A differentiate that set of fossils from Group B with the Taung child. Some characters in Group A are exhibited by the Makapansgat fossils MLD 1 (type specimen of *A. prometheus*) and MLD 2, a juvenile mandible with large *Paranthropus*-like cheek teeth (Dart, 1948a,b). It therefore seemed most appropriate to place the Group A fossils into an already named species, rather than create a new one.

The StW 573 cranium can be compared with the fossils of these two groups in order to determine its closest affinities. It matches Group A, i.e., *A. prometheus*, in all features except cheek teeth size, and that is discussed below.

Based on the above observations, the following species definitions are offered:

### *Australopithecus africanus* (Dart 1925)

A species of the genus *Australopithecus* characterised by the following features: relatively small face and small braincase with cranial capacity of less than 500 cc; sides of braincase sloping superomedially; high convex frontal squame; presence of a metopic ridge; moderate convergence of temporal lines with no sagittal cresting; narrow interorbital distance; convex midface; gracile zygomatic arch with reduced masseteric attachment; formation of a shelf over the external auditory meatus; well-defined separation between clivus and nasal floor, with no nasal guttering; vertically oriented incisors; relatively uncorrugated clivus; relatively small canines; absence of diastema; small round-sectioned upper canine roots; anteriorly deep, relatively short palate; cheek teeth with vertical sides and high pointed cusps; more generalised tooth wear in adults; relatively small mandible with thick corpus and thickened base; low occipitomastoid crests.

### *Australopithecus prometheus* (Dart 1948)

Large face and braincase with cranial capacity above 500 cc; vertical sides to braincase with prominent parietal bossing; low concave frontal squame; no metopic ridge; strongly converging temporal lines that often unite in a sagittal crest; flat midface; broad interorbital distance; robust zygomatic arch with deep malar and large masseteric attachment; no shelf over external auditory meatus; inferior nasal margin smoothly concave; procumbent upper incisors and roots; clivus corrugated by incisor roots; large canines; large flattened upper canine roots; shallow anterior palate; long rectangular palate; large upper incisor-canine diastema; large cheek teeth; side of cheek teeth slope towards occlusal surface; low bulbous cheek teeth cusps; excessive anterior tooth wear in adults; large mandible with big teeth but relatively slender corpus; prominent occipitomastoid crests.

A third species that requires some clarification is *Homo habilis*, as there has been disagreement over which specimens belong to that taxon, and one in particular, i.e., StW 53, is relevant to the identification of Sterkfontein *Australopithecus* (Leakey et al., 1964; Leakey et al., 1971; Leakey, 1974;

Hughes and Tobias, 1977; Walker, 1981; Johanson et al., 1987; Tobias, 1991; Clarke, 2013, 2017). The type specimen of *Homo habilis* was the OH 7 parietals and mandible from Olduvai Gorge, Bed I, and that certainly seems to have had a larger brain than *A. africanus*. However, other specimens from Olduvai, such as OH 24 (Leakey et al., 1971), strongly match fossils of *A. africanus* and differ dramatically in both brain size and facial structure from KNM ER 1470, which has parietals and brain size matching those of the type specimen OH 7. A detailed analysis of the dispute over these fossils is provided in Clarke (2017). The StW 53 cranium was classed as early *Homo* by Hughes and Tobias (1977) because they considered it came from a stone tool-bearing deposit. However, the morphology of that specimen, which Tobias eventually classed as *Homo habilis* (Tobias, 1991), matches very closely that of the first adult *Australopithecus* cranium from Sterkfontein, TM 1511 (Fig. 25), just as the OH 24 morphology matches very closely that of *Australopithecus* cranium Sts 17 (Fig. 26). Furthermore, the StW 53 cranium did not come from the tool-bearing Member 5, but from a hanging remnant of Member 4 that contains no artefacts (Kuman and Clarke, 2000). There is no justification morphologically for separating either OH 24 or StW 53 from *A. africanus*, and thus they are considered here as representatives of that species. Another Olduvai Bed I specimen, OH 62, was classed as *Homo habilis* because of its resemblance to StW 53 (Johanson et al., 1987). That too seems most probably to be an East African representative of *A. africanus.* There is, however, one Olduvai Bed I specimen, OH 65, that does differ dramatically in morphology from *Australopithecus* and which resembles *Homo habilis* in the form of the 1470 cranium (Clarke, 2012). A South African specimen which resembles OH 65 is SK 27 from Swartkrans (Clarke, 2017).

**Fig. 25.**
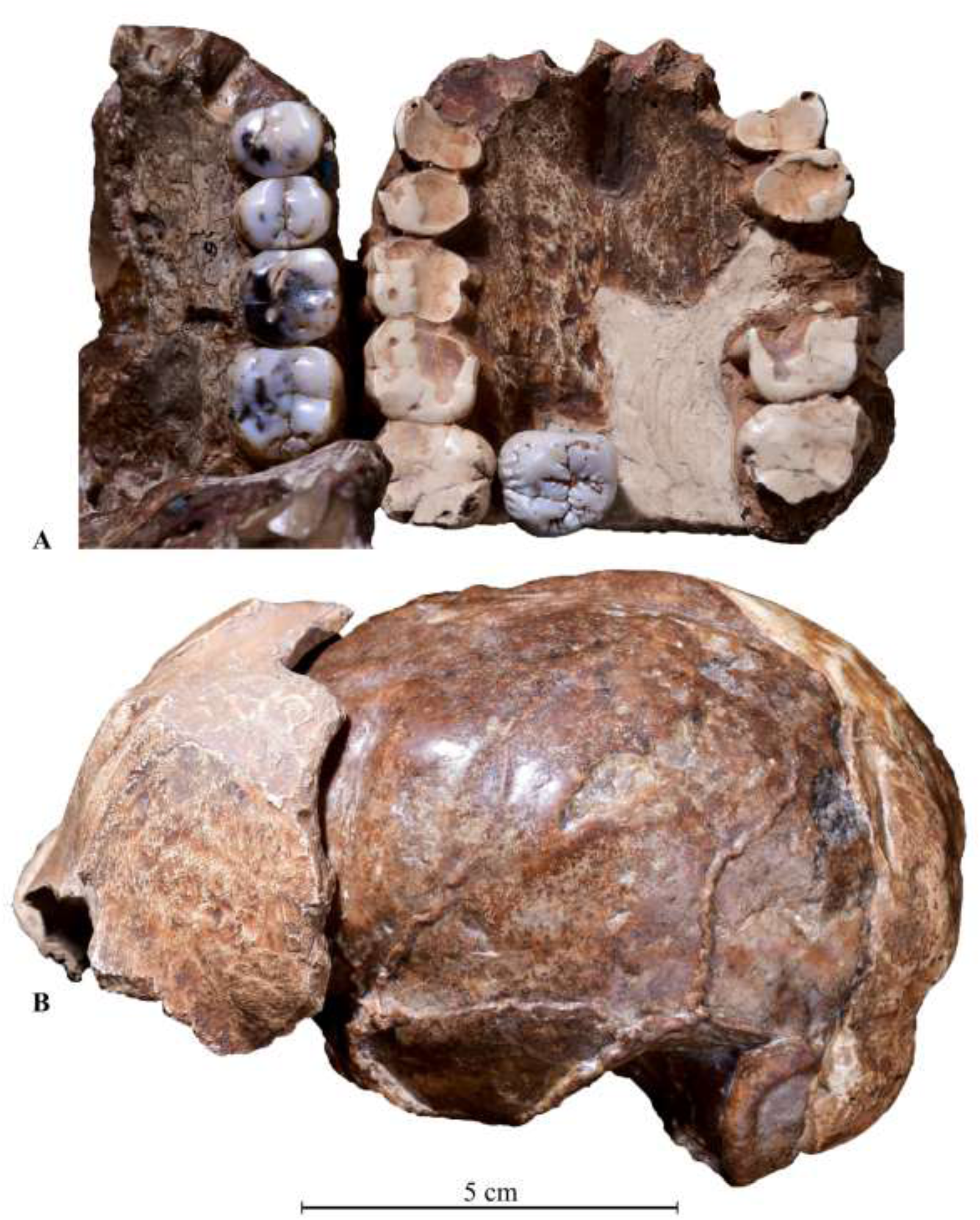
The close similarity of the TM 1511 palate and dentition with that of StW 53. A) on left is the TM 1511 palate with P^3^ to M^3^. The latter tooth is isolated and placed on the StW 53 palate (right). B) The close fit of the StW 53 frontal bone onto the frontal lobe of the TM 1511 (Sts 60) endocranial cast.

**Fig. 26.**
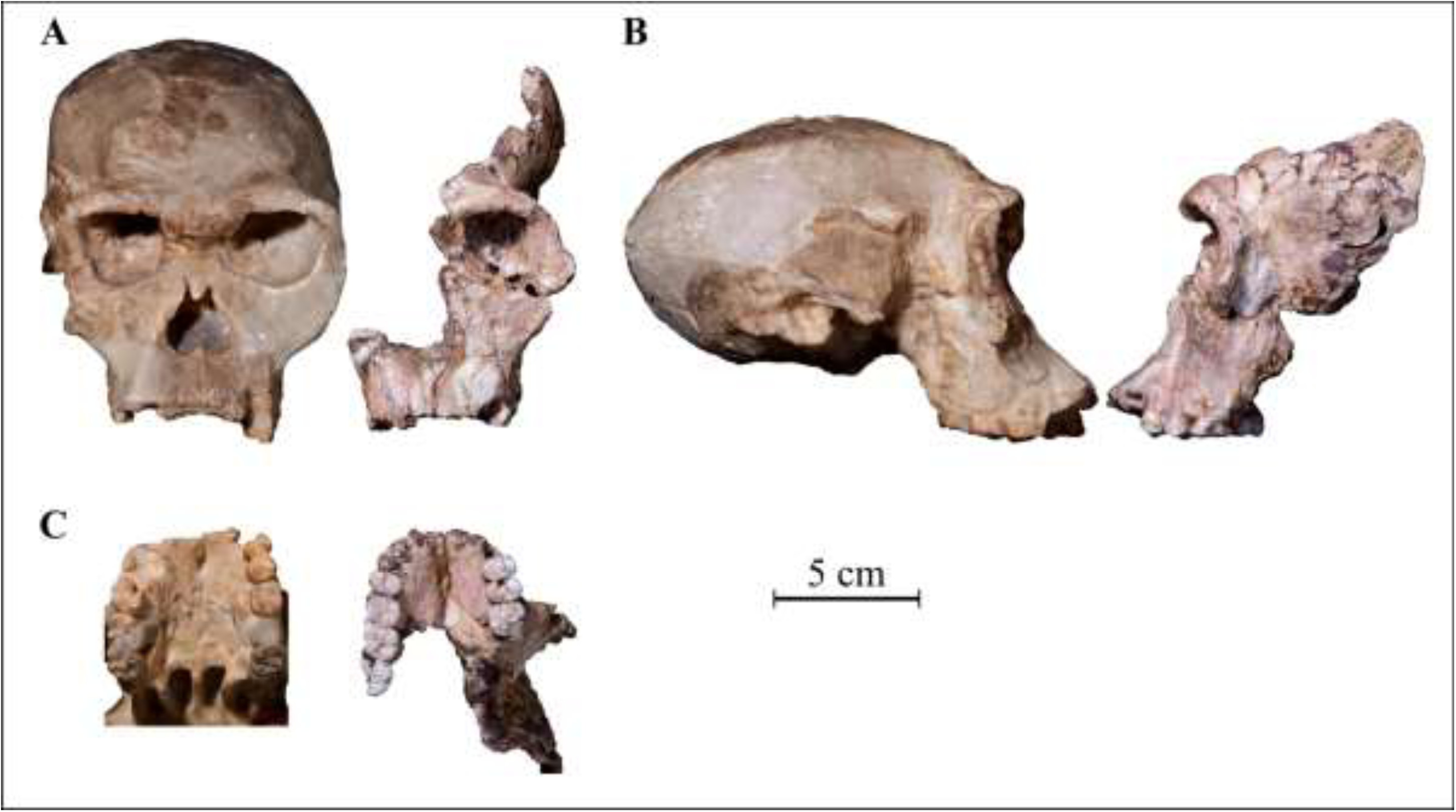
Comparisons of the OH 24 cranium and Sts 17 crania. A) facial view with OH 24 at left and Sts 17 at right; B) profiles views of OH 24 (left and Sts 17 (right); C) palatal views of OH 24 (left and Sts 17 (right).

The following definition can be offered for *Homo habilis*:

### *Homo habilis* (Leakey, Tobias and Napier 1964)

A species of the genus *Homo* characterised by the following features: large braincase with cranial capacity up to 750 cc; laterally expanded parietals; sides of braincase vertical with parietal bosses; large face with relatively broad and flat midface; high convex frontal squame; broad frontal with widely separated temporal lines anteriorly and posteriorly; no metopic ridge; prominent glabella; medium interorbital distance; anteriorly prominent nasal skeleton; sharply defined lower nasal margin with nasal sill; deep malar; relatively flat and broad lower face; wide orthognathic nasoalveolar clivus; large broad and deep palate; P3 roots situated posterolateral to the canine roots; no diastema present; moderately sized cheek teeth with large canines and incisors; mandible of moderate size but with robust corpus; no premaxillary suture on the facial surface.

## 6. Discussion

The total morphological pattern of StW 573 aligns it with certain Makapansgat and Sterkfontein fossils that, as a group, exhibit numerous characters that distinguish them from a second morphological group consisting of fossils from Taung, Sterkfontein and Makapansgat. This first group includes MLD 1, the type specimen of *A. prometheus*, and is thus considered to represent that species. The second group includes the Taung child, the type specimen of *A. africanus*, and the group as a whole is therefore considered to show characters representing that species. Both of these species were contemporary and found in the same cave breccias in both Sterkfontein Member 4 and Makapansgat.

Previously, similarities or differences between South African *Australopithecus* fossils had to be determined on fragments or on teeth, partial crania and mandibles. The advantage of StW 573 is that it is a complete skull against which more fragmentary fossils can be compared or contrasted. Thus this skull provides a focal point for the clustering of fossils exhibiting the same characters. Furthermore the fact that this skull of an apparent *A. prometheus* belongs with an almost complete skeleton means that isolated limb bones and fragments can be compared or contrasted and more confidently assigned to one or the other species. When the sample of Sterkfontein, Makapansgat and Taung *Australopithecus* fossils is divided between the two species of *A. africanus* and *A. prometheus*, it is possible to interpret certain size and robusticity variation within each group as probably representing sexual dimorphism. Thus, for *A. africanus*, Sts 5 with a smaller canine socket and narrower cranial base than StW 53 is apparently female, whilst StW 53 is apparently male. Sts 52 seems to be male, whilst TM 1512 with similar facial morphology but smaller teeth seems to be female. In the other group, i.e., *A. prometheus*, StW 505 in its great size and robusticity is surely a male, contrasting with the smaller cranium of Sts 71, which is probably female. The young individual StW 252 with very large cheek teeth and anterior dentition, as well as large braincase, can be considered male. The StW 573 skull, having small braincase, narrow cranial base, and anterior dentition which, although large is smaller than that of StW 252, appears to be female.

The cheek teeth of StW 573 are smaller than one would expect for a female of *A. prometheus*, considerably smaller than the female Sts 71 (see Table 2). However, visual inspection of the exposed sides of the upper and lower third molars does show some bulging of the sides typical of *A. prometheus*. The small cheek teeth size seems to be beyond normal variation and could possibly relate to the earlier geological age of StW 573. It has been dated to ca 3.67 Ma (Granger et al., 2015) and derives from an older cave infill than Member 4, which contains the other *A. prometheus* fossils at Sterkfontein. Extreme megadontia found in some *Australopithecus* and *Paranthropus* fossils seems to be a derived condition. It is possible then that the smaller cheek teeth in the 3.67 Ma StW 573 represent an ancestral condition from which the later megadont *A. prometheus* derived. This aspect would benefit from further investigation and an enlarged sample of *A. prometheus* fossils from that earlier time period.

In describing his reconstruction of the Sterkfontein StW 252 cranium, Clarke (1988) suggested that it was ideally placed temporally and morphologically to be a member of a species ancestral to *Paranthropus*. It had retained primitive ape-like characters of prognathism with large incisors and canines, but it had the *Paranthropus*-like features of enlarged, inflated cheek teeth, together with midfacial and supraglabellar hollowing. He noted that temporally it was ‘situated between an ancestral *Australopithecus*, possibly *A. afarensis*, and the earliest *Paranthropus*.’ The discovery and dating of StW 573 now indicate that an early form of *A. prometheus* was contemporary with early *A. afarensis* at Woranso Mille, Ethiopia and Laetoli, Tanzania (both ca 3.6 Ma). Hence it could not have descended from *A. afarensis* but both of them came from yet earlier ancestry, of which *A. anamensis* at 4.2 Ma seems to be one good example. The position of *A. prometheus* temporally in relation to *Paranthropus* depends upon the dating of Sterkfontein Member 4, which is currently under renewed investigation. Currently there is no reason to suppose that Member 4 is younger than 2.5 Ma and it might well be considerably older. The faunal age estimate for that member is between 2.4 to 2.8 Ma (Vrba, 1985). Debate over a younger age for Member 4 has in particular been generated by palaeomagnetic results on flowstones, but it is now known that these are younger post-depositional calcites that formed in voids created by subsidence of breccias (see Bruxelles et al. 2018, this issue, for further detail). Inspired by a preliminary result that suggests a greater age, D. Granger is currently conducting more extensive dating with the cosmogenic isochron method to better determine the age range of Member 4.

Thus there still remains a possibility, subject to dating verification, that *A. prometheus* was ancestral to *Paranthropus robustus*, and that both of them were contemporary with the evolving lineages of *A. africanus* and early *Homo* (Leakey, 1974; Walker, 1981; Clarke, 2017). In East Africa at Olduvai Gorge Bed I, Tanzania, a late form of *A. africanus* (OH 24 and OH 62--Clarke, 2012) was contemporary with both *Homo habilis* (OH 7 and OH 65) and *Paranthropus boisei* (OH 5). The *Homo* lineage is also now considered to go back at least 2.8 million years, as indicated by the fossil mandible from Ledi Geraru, Ethiopia (Villmoare et al., 2015).

One East African fossil that does have some resemblance to *A. prometheus* is the partial cranium (Bou-VP-12/130) from Bouri, Ethiopia (Asfaw et al.,1999). This is the type specimen of *Australopithecus garhi* and dates to 2.5 Ma. It is similar in overall shape of calotte and maxilla, with medially encroaching temporal lines on the frontal enclosing a depressed postglabellar squame, a sagittal crest, convex clivus with procumbent incisors, and relatively large canines with very large cheek teeth. However, Asfaw et al. (1999) state that ‘in cranial anatomy it is definitely not *A. africanus.*’ It should be noted that they, like most other researchers, lumped all fossils from Sterkfontein and Makapansgat with Taung into *A. africanus*. Hence they were implying that its cranial anatomy was not even like those fossils which are here considered as belonging to *A. prometheus*. *A. garhi* certainly does differ from *A. prometheus* in the wear pattern of its teeth, which shows lingual wear on the P^4^ and M^1^ but hardly any wear on the anterior dentition, even though it is an adult with fully erupted third molars. Another apparent difference is in the brain size. Asfaw et al. (1999) concluded that the cranium is male and stated that Holloway had determined a cranial capacity of about 450 cc. Even allowing for individual variation, this brain size of an apparent male does seem considerably smaller than Holloway’s (2012) estimate of slightly above 575 cc for the StW 505 male *A. prometheus*. As earlier mentioned, the cranial capacity for StW 573 is even smaller than Bouri, but it is an apparent female and is more than one million years older.

The skull of StW 573 from the Sterkfontein Caves Member 2 displays the greatest similarity in the sum of its morphological characters with fossils from Member 4 Sterkfontein and Member 3 Makapansgat that belong to the species *Australopithecus prometheus* (see Fig. 27 for skull comparisons). The only notable difference is in the small size of its cheek teeth, and that may be related to its earlier geological age of ca 3.67 Ma. StW 573 differs from *Paranthropus* in having large canines and incisors, which are very heavily worn. This contrasts with the derived condition of reduction of those teeth in *Paranthropus*, which are in marked contrast to their enlarged premolars and molars.

**Fig. 27.**
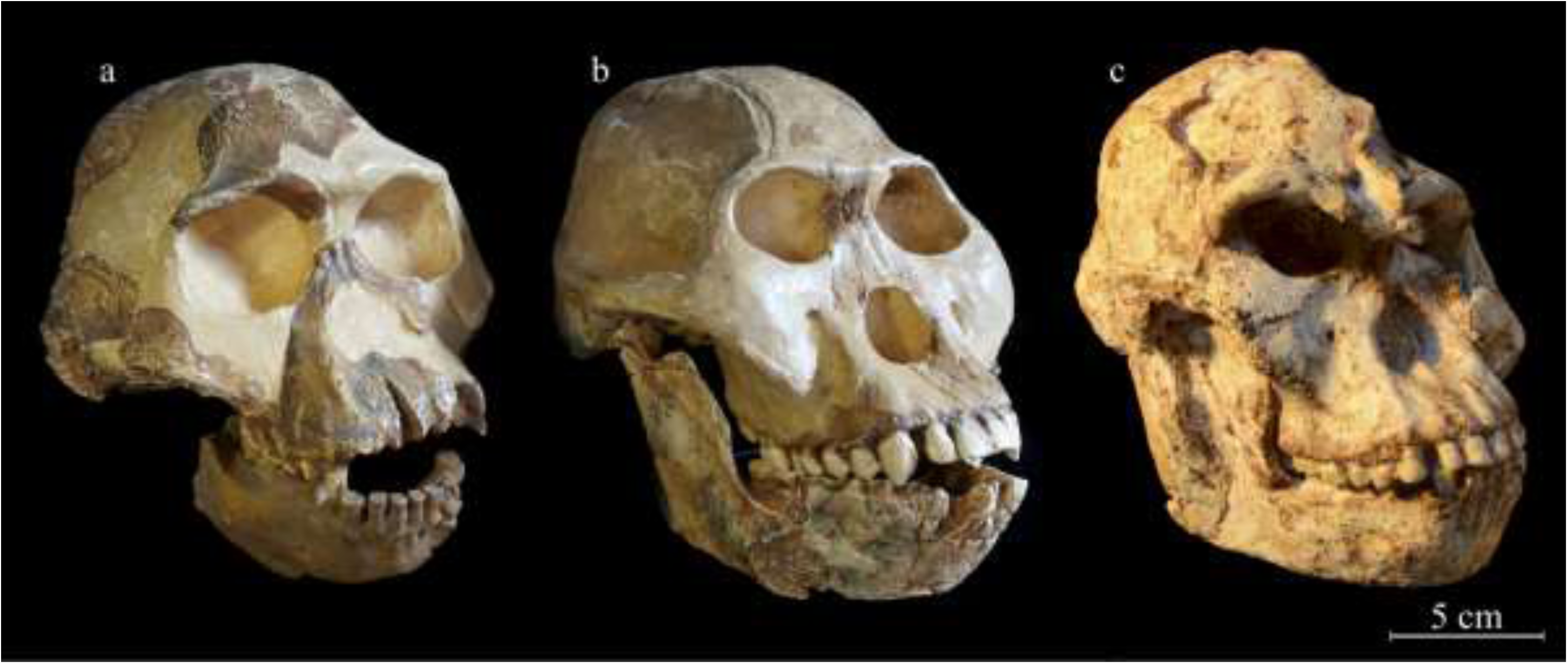
Comparison of *A. africanus* and *A. prometheus* skulls. a) StW 53 male cranium articulated with MLD 18 mandible, both *A. africanus*. b) StW 252 young male cranium articulated with Sts 36 mandible, both *A. prometheus*; c) StW 573 female *A. prometheus* complete skull. Note the much larger skulls, flatter faces, and flatter frontal bones of both male and female *A. prometheus*, contrasting with the smaller size and convex midface and frontal bone of the male *A. africanus*.

It has been demonstrated in the previous comparative section that the anatomical characters which align StW 573 with *A. prometheus* also distinguish it from *A. africanus*. These distinctions can be seen also between StW 573 and South African cranial fossils that have been assigned to a new species, *A. sediba* (de Ruiter et al., 2013). Not only is StW 573 considerably older geochronologically and much larger in size, but *A. sediba* has more in common with *A. africanus* than with *A. prometheus*. De Ruiter et al. (2013) state, ‘The closest morphological comparison to *A. sediba* within the australopiths is *A. africanus*, as the two share numerous similarities in the cranium, face, palate, mandible, and teeth.’ Kimbel and Rak (2017) state that there is ‘clear evidence that *A. sediba* was uniquely related to *A. africanus*.’ Both De Ruiter et al. and Kimbel and Rak include all Sterkfontein Member 4 specimens as *A. africanus*. Thus they are mixing together distinct characters of the two species, *A. prometheus* and *A. africanus*, which we here and in previous studies (Clarke 1988, 1994, 2013) argue should not be lumped together. Kimbel and Rak (2017) further observe that ‘key aspects of MH1’s resemblance to *Homo* can be accounted for by its immaturity.’ In addition to that point, we here note some features listed by de Ruiter et al. (2013) in their comparative table that give a misleading characterization of the differences between *A. sediba* and *A. africanus*.

These are:

1) The cranial capacity, which they list as ‘small’ in *A. africanus*, but they include in that taxon the specimen StW 505 (which is here presented as *A. prometheus*). As earlier noted, Holloway (2012) gave the cranial capacity of StW 505 as greater than 575 cc. That is considerably larger than cranial capacities for *A. africanus* and *P. robustus*. Even allowing for its immature development, the cranial capacity of MH1 at 420 cc (assessed as 97% of its brain growth--de Ruiter et al., 2013) is very much smaller than 575 cc.

2) De Ruiter et al. (2013) list incisor procumbency as being variable in *A. africanus*, contrasting with vertical incisors in *A. sediba*. In fact, as discussed in this paper, procumbent incisors are characteristic of *A. prometheus*, whilst many other Member 4 fossils, which we differentiate as *A. africanus*, have vertically placed incisors. Thus in this feature, *A. sediba* aligns with *A. africanus* and not with *A. prometheus*.

3) De Ruiter et al. (2013) list ‘derived facial morphology of robust australopiths’ as being absent in *A. africanus*. This is certainly not correct in their taxonomic hypodigm of the species which includes fossils which we attribute to *A. prometheus*. Clarke (1988, 1994) discussed several features of the face and frontal that were *Paranthropus*-like in StW 252 and Sts 71, which, together with other specimens now including StW 505, are considered diagnostic of *A. prometheus*.

4) De Ruiter et al. (2013) list post-canine megadonty as being absent in *A. africanus*, in which category they include specimens such as StW 252, StW 384, and StW 498 that we regard as representing *A. prometheus* because of their post-canine megadonty and the inflated *Paranthropus*- like morphology of their cusps. In fact, specimen StW 384 has post-canine teeth equal in size to those of the large Peninj mandible of *P. boisei*, as illustrated in Clarke (1994). Several specimens of *A. prometheus* have post-canine teeth which are even larger than the post-canine teeth of some *P. robustus* individuals, e.g., the StW 252 *A. prometheus* cheek teeth are larger than the SK 46 large male *Paranthropus* cheek teeth (Clarke, 1988).

Thus *A. sediba* does not align with *A. prometheus*, including StW 573, in its cranial and dental morphology, but it does have characteristics in common with *A. africanus* if the fossils of *A. prometheus* are separated from that taxon.

## 7. Conclusion

The StW 573 skull is complete, belongs to a mature individual, and belongs with a skeleton which is the only *Australopithecus* fossil so far discovered in the Member 2 breccia of Sterkfontein Caves. It is stratigraphically older than the large number of *Australopithecus* fossils from Member 4 of the Sterkfontein Caves and has been dated by cosmogenic burial dating to ca 3.67 Ma, making it a contemporary of early *A. afarensis* fossils from East Africa (Woranso Mille and Laetoli). Although StW 573 has some morphological features similar to *A. afarensis*, its closest morphological counterparts are some of the *Australopithecus* fossils from Sterkfontein and Makapansgat, including StW 505, StW 252, StW 183, StW 498, Sts 71, Sts 36, Sts 7, MLD 2, MLD 9, TM 1516, and the type specimen of *A. prometheus* MLD 1. These show a total morphological pattern that differentiates them from the total morphological pattern exhibited by a cluster of other Sterkfontein and Makapansgat fossils, including TM 1511, Sts 5, StW 53, Sts 17, Sts 19, Sts 52, TM 1512, StW 404, MLD 6, MLD 18, MLD 40, and the type specimen of *A. africanus*, the Taung child.

Although the type specimen of *A. prometheus* is only the back part of a braincase, it does show features linking it to the StW 252 and StW 505 group. These features are the vertical sides to the parietals curving around parietal bosses that produce a wide upper braincase, the large estimated cranial capacity of 500 cc, and the indications from the closeness of the temporal lines that it may have had a sagittal crest.

As StW 573 has strong morphological similarities to StW 505 and StW 252, which in turn are similar to the MLD 1 *A. prometheus* type specimen, it seems most appropriate to assign StW 573 to *A. prometheus*. There are some differences in the nuchal area of StW 573 which has a pronounced nuchal crest, a pronounced inion, and very prominent occipitomastoid crests. In MLD 1, the nuchal crest and inion are only slightly marked and there are no indications of occipitomastoid cresting. These differences, however, do not seem sufficient to separate MLD 1 taxonomically from StW 252, StW 505, or StW 573 because the general shape of the braincase does not align it with Sts 5 or StW 53.

It could be argued that the more rugged nuchal muscle attachments on StW 573 indicate that it is male, but certain other features such as the narrowness of the cranial base, the small mastoid processes, and the smaller canines and incisors in comparison to those of StW 252 suggest that it is female. Such a conclusion is supported by the greater sciatic notch of the pelvis, which is more open than the narrowed sciatic notch in StW 431 which appears to be male.

Rather than creating yet another new species for StW 573, it seems most appropriate for the present to assign it to the existing species *A. prometheus*, with which it aligns more closely than with any other *Australopithecus* species. Those who may find it difficult to accept that two species of *Australopithecus* (*A. africanus* and *A. prometheus*) could occupy the same areas at the same time should note that at both Makapansgat and Sterkfontein two species of monkeys, i.e., *Parapapio jonesi* and *Parapapio broomi*, are abundantly represented, and there has never been any question about their taxonomic separation. Furthermore, among carnivores, the leopard *Panthera pardus* and the lion *Panthera leo* occupy the same territories.

Fossils of *Paranthropus* and early *Homo*, whilst not of one genus, do illustrate that early human relatives of different species do occupy the same territories at the same times in both East and South Africa. Their dental and cranial differences suggest that they fed differently and thus would not have necessarily been competitors for food resources. The same could have been true of *Australopithecus africanus* and *Australopithecus prometheus*, as the latter has a more *Paranthropus*- like cheek dentition and cranial structure, as well as a different pattern of tooth wear. Elton (2001) discussed the sympatry of *Parapapio jonesi* and *Parapapio broomi* at Sterkfontein, referring to Fleagle (1988) on the sympatry of modern *Cercopithecus* species that forage in mixed species groups in West African forest habitats. She also noted in Asia that the secondary forest-dwelling *Macaca fascicularis* is more arboreal than the sympatric *Macaca nemestrina* that lives in upland and hilly areas and often travels on the ground (Fleagle, 1988).

## Acknowledgments

The excavations that resulted in the recovering of this skeleton were carried out under permits from the South African Heritage Resources Agency. We are grateful to PAST (the Palaeontological Scientific Trust, South Africa) for major financial support and advocacy, without which this project could not have been completed; Andrea Leenen, Rob Blumenschine, PAST founding member John Cruise, and the late Dusty van Rooyen have in particular provided valuable logistical support over the years. The following donors have provided generous financial assistance over two decades: Standard Bank and J.P. Morgan via PAST, the DeBeers Chairman’s Fund, the National Research Foundation of South Africa (NRF), the Department of Science and Technology (DST, formerly DACST), the Wenner Gren Foundation, the L.S.B. Leakey Foundation, the Ford Foundation, the Mott Foundation, the Embassy of France in South Africa, and the National Geographic Society. Phillip Tobias and Alun Hughes introduced us to Sterkfontein research. We appreciate the dedication of Stephen Motsumi, Nkwane Molefe, Abel Molepolle, and Andrew Phaswana to the excavation, preparation and casting work on the skeleton, as well as the hard work of the entire Sterkfontein field crew. Paul Myburgh devoted much time and his own resources over 20 years to meticulous filming of the excavation and preparation process. KK wishes to acknowledge her NRF research grants (Nos. 82591 and 82611) and university research incentive funds. Dominic Stratford is the current director of Sterkfontein research since R.J.C’s retirement and has provided major funding for the work and support of the technical staff in the last several years for which we are grateful (Stratford NRF grant number 98808). We greatly appreciate the many years of dedicated research contributed by our two colleagues, Travis Pickering and Jason Heaton, who have for many years worked assiduously on many Sterkfontein hominids and have had a long-term involvement in the the Member 2 research. We are further grateful to all the members of the Little Foot academic team for their collaborative work and generous sharing of data and images: Travis Pickering, Jason Heaton, Dominic Stratford, Robin Crompton, Amelie Beaudet, Laurent Bruxelles, Kristian Carlson, Tea Jashashvili, and Juliet McClymont, who are all co-authors on papers in this special issue. We thank Matt Lotter for his excellent photography and assembling of figures. We wish also to acknowledge our dating, karst stratigraphy and laser scanning colleagues for their efforts over these years: Darryl Granger, Ryan Gibbon, Marc Caffee, Tim Partridge, Richard Maire, Richard Ortega, Jose Braga and Gerard Subsol. The collaboration of all these colleagues has been invaluable. Finally, it is Sarah Elton who has made this special issue entirely possible. She has our deep respect and gratitude, as do Eric Delson and three other anonymous reviewers who have helped improve this paper.

